# Three-dimensional imaging of vascular development in the mouse epididymis: a prerequisite to better understand the post-testicular immune context of spermatozoa

**DOI:** 10.1101/2022.08.26.505406

**Authors:** Christelle Damon-Soubeyrand, Antonino Bongiovanni, Areski Chorfa, Chantal Goubely, Nelly Pirot, Luc Pardanaud, Laurence Pibouin-Fragner, Caroline Vachias, Stéphanie Bravard, Rachel Guiton, Jean-Léon Thomas, Fabrice Saez, Ayhan Kocer, Meryem Tardivel, Joël R. Drevet, Joelle Henry-Berger

## Abstract

Long considered an accessory tubule of the male reproductive system, the epididymis is proving to be a key determinant of male fertility. In addition to its secretory role in ensuring functional maturation and survival of spermatozoa, the epididymis has a complex immune function. Indeed, it must manage both peripheral tolerance to sperm antigens foreign to the immune system and the protection of spermatozoa as well as the organ itself against pathogens ascending the epididymal tubule. Although our knowledge of the immunobiology of this organ is beginning to accumulate at the molecular and cellular levels, the organization of blood and lymphatic networks of this tissue, important players in the immune response, remains largely unknown. In the present report, we have taken advantage of a VEGFR3:YFP transgenic mouse model. Using high-resolution three-dimensional (3D) imaging and organ clearing coupled with multiplex immunodetections of lymphatic (LYVE1, PDPN, PROX1) and/or blood (PLVAP/Meca32) markers, we provide for the first time a simultaneous deep 3D view of the lymphatic and blood epididymal vasculature in the mature adult mouse as well as during postnatal development.

**Graphical abstract:** Summary of the expansion of the conventional and hybrid lymphatic vasculature during postnatal development of the murine epididymis.

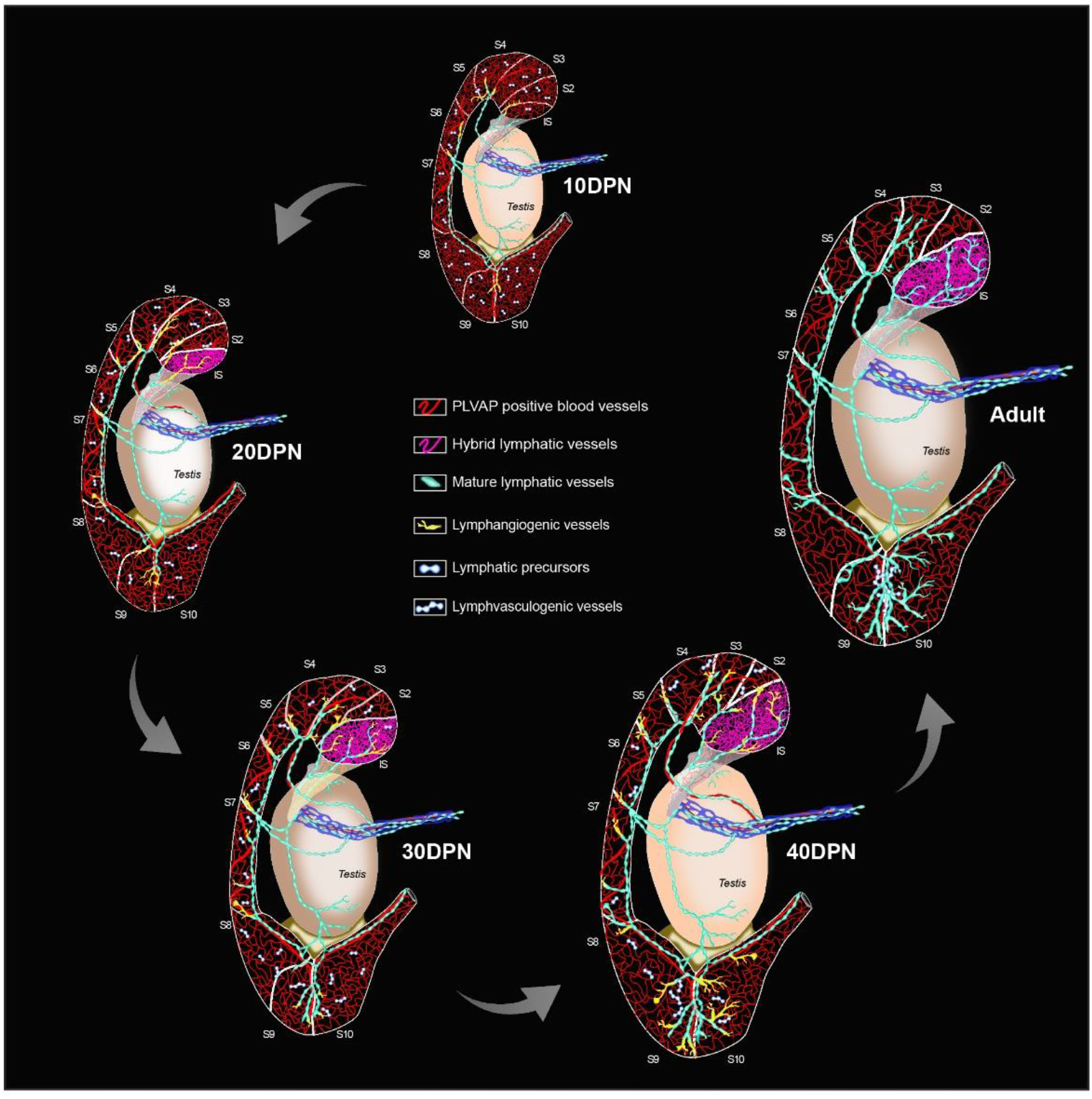

## Introduction

Studies regarding the etiology of male infertility have revealed that at least 15% of cases have a clear immunological origin, a share that is most likely underestimated if idiopathic infertilities were to be included (Dohle et al., 2005; Jungwirth et al 2012). In the male, they may be the result of chronic inflammatory situations, acute bacterial and/or viral infections, or autoimmune processes along the genital tract leading to the production of anti-sperm antibodies (ASA; for a recent review, see Shibahara et al., 2021). These situations are difficult to diagnose and treat, in part because our knowledge of immune responses in male accessory organs is limited.

Unlike the testis, where mammalian evolution has chosen to tightly seal the germline from the immune system—conferring immune privilege to this tissue (Meinhardt and Hedger, 2011)—the epididymis tubule faces multiple immune challenges (Guiton et al., 2013; Hedger, 2011). On the one hand, the epididymis tubule is the testis gatekeeper, preventing ascending pathogens from reaching the immune-privileged seminiferous tubules. To do so, it should be endowed with all the power of a full and efficient immune response toward exogeneous antigens. On the other hand, the epididymis must maintain self-tolerance toward spermatic antigens that are unique to this cell lineage and appear at puberty long after the establishment of the self-immune repertoire (Goodnow, 1996). Understanding how this dual action of peripheral tolerance toward sperm antigens (Mueller et al., 2010) versus efficient immune survey toward non-self-antigens is performed is of particular interest for infertility issues, but also more largely in other settings where this situation occurs.

This has prompted recent research aiming at identifying immune cells and molecules that could participate in the finely orchestrated epididymal immune physiology. This research has led to the characterization of a dense network of peritubular antigen-presenting cells [APC] (Da Silva et al., 2011), interstitial and intra-epithelial lymphocytes of distinct sub-families (Voisin et al., 2018), as well as of immunosuppressive players such as transforming growth factor beta [TGF-β] (Pierucci-Alves et al., 2018; Voisin et al., 2020) and indoleamine/Tryptophane dioxygenase activity [IDO/TDO] (Britan et al., 2006; Jrad-Lamine et al., 2011, 2013).

Because the immune picture of the mammalian epididymis would not be complete without a clear view of the blood and lymphatic circulating networks—which are crucial in managing immune responses—we choose to explore them in the mouse. This approach was necessary, as existing data are rather old and scarce, especially for the lymphatics (Abe et al., 1984; Hirai et al., 2010; Itoh et al., 1998; McDonald and Scothome, 1988; Perez-Clavier et al., 1982; Suzuki, 1982). Characterization of the lymphatic vasculature in mammals is tricky because it shares common embryonic origins with the blood vessels (Baluk et al., 2008; Srinivasan et al., 2007). Nevertheless, a step forward was made with the identification of lymphangiogenic factors (vascular endothelial growth factor [VEGF-C and VEGF-D]) and their receptor (VEGFR3), which are now considered the major markers of lymphatics (Kaipainen et al., 1995; Mäkinen et al., 2001). To ensure confidence in the identification of lymphatic structures, one should use of a combination of markers, including VEGFR3, the lymphatic vessel endothelial hyaluronan receptor 1 (LYVE1), the Prospero-related homeobox 1 (PROX1) transcription factor, and podoplanin (PDPN) (Banerji et al., 1999; Breteneder-Geleff et al., 1999; Hamrah et al., 2003; Kivelä et al., 2016; Mouta-Carreira et al., 2001; Partanen et al., 2000; Wigle et al., 1999). We have followed this approach in the present study, using three lymphatic markers together with plasmalemma vesicle-associated protein (PLVAP/MECA32) to specifically identify fenestrated blood vessels (Stan et al., 2004). In addition, we have taken advantage of a transgenic mouse model that expresses the fluorescent reporter YFP under the control of the VEGFR3 promoter (Calvo et al., 2011). Using the power of three-dimensional (3D) reconstruction of high-resolution imaging with light-sheet microscopy after epididymis 3DISCO clearing (Belle et al., 2014; Ertük et al., 2012), we present here an in-depth analysis of both the blood and lymphatic networks of the epididymis in the adult mouse as well as during postnatal ontogenesis.

## Results

### An abundant lymphatic network drains the epididymis

To gain insight into the lymphatic network of the mouse epididymis, we used the VEGFR3:YFP mouse model. Figure 1 shows representative *in toto* views of the lymphatic vascularization of the testis and epididymis of an adult (A) and 10 days postnatal (DPN) mouse (B and C).

**Figure 1:**
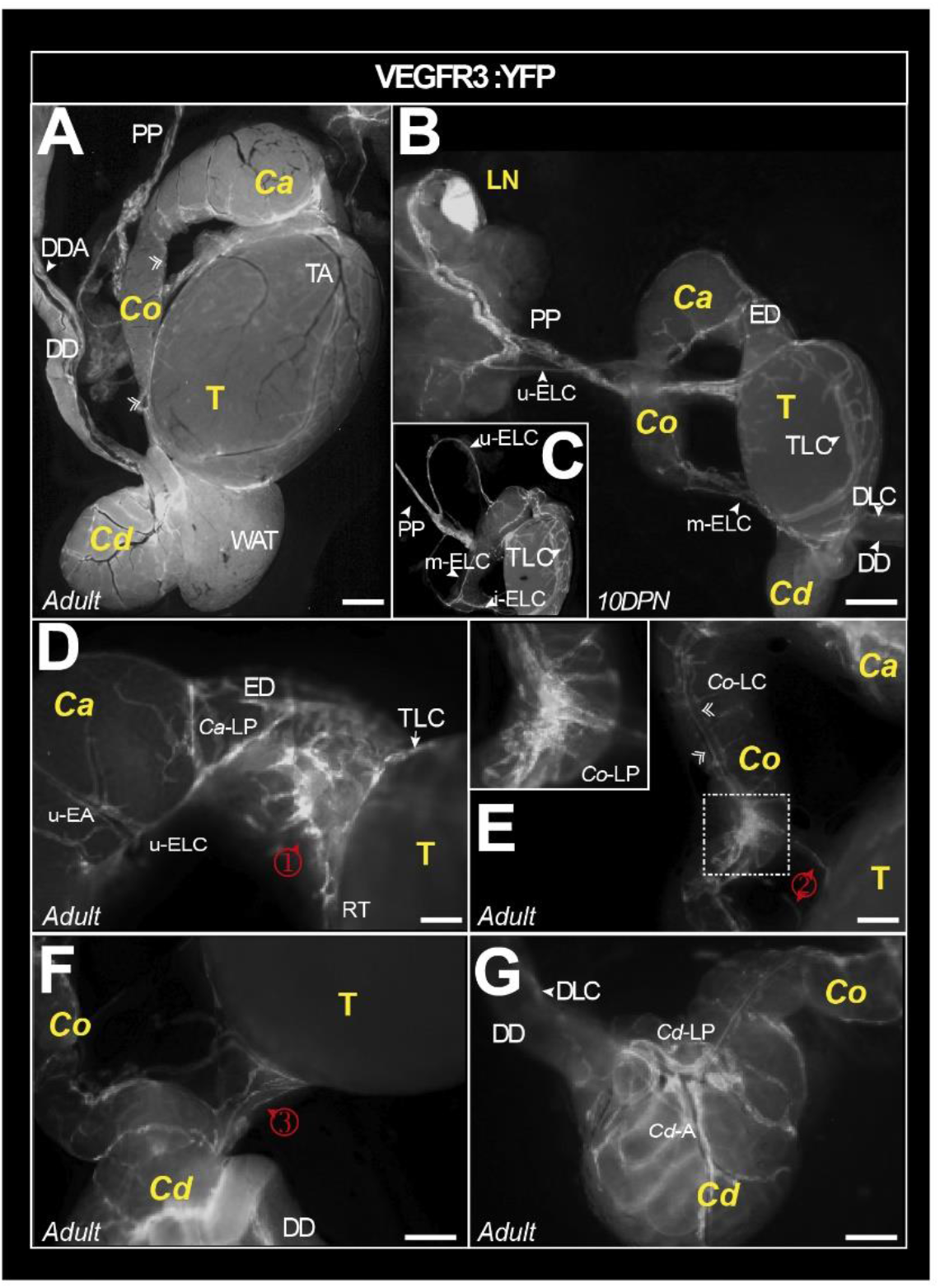
Macroscopic view of the lymphatic vasculature of the epididymis and testis in the VEGFR3:YFP model. Representative image of the lymphatic vasculature of the adult epididymis and testis observed with a Leica binocular loupe (A). VEGFR3:YFP lymphatics arc visible in the caput, corpus, and cauda regions. Lymphatics responsible for epididymal and testicular drainage follow the pampiniform plexus (PP) before reaching a lymph node (B). The superior and lateral epididymal lymphatic collectors (upperELC and medianELC, respectively) drain lymph from the caput and corpus, respectively, before joining the main testicular lymphatic collector at the PP. The latter is connected to a main collector that surrounds the testis (arrowhead) and branches into a rich network (A-C). There arc numerous lymphatic connections between the lymphatics of the epididymis and the main lymphatic collector of the testis, notably through a lymphatic network at the level of the efferent ducts (① in D), corpus (② in E), and cauda (③ in F). Through the capsule covering the epididymal duct, there is also fluorescence that outlines the tubules, weakly in the caput (D) and more intense in the cauda (F and G). The scale bar corresponds to 1 mm. AT =adipose tissue; *Ca* = *caput*;, *Cd* = *cauda*, *Co* = *corpus*, DD =deferent duct; DDA =deferent duct artery; DLC = deferent lymphatic collector; DPN = days postnatal; ED = efferent duct; LELC = lower epididymal lymphatic collector; LN; lymph node; SEA = superior epididymal artery; SELC =superior epididymal lymphatic collector; T =testis; TA =testis artery; TLC =testicular lymphatic collector; *Cd*-A = Caudal artery; *Cd*-LP = Caudal Lymphatic Plexus. ① = caput-testis lymphatic connection; ② = corpus-testis lymphatic connection; ③ = cauda-testis lymphatic connections between testis and epididymis.

There is intense vascularization at the epididymis–testis interface regardless of the level of the epididymis (*caput, corpus*, or *cauda*). Figure 1B shows that the lymphatic drainage joins the main testicular lymphatic trunk at the *pampiniform plexus* (PP) and follows it to a highly fluorescent structure that corresponds to the most proximal lymph node (LN). Inset 1C and Figure 1D show the connections of the collectors emanating from the *caput* (upper epididymal lymphatic collector [u-ELC] in C and D) and the *corpus* (median epididymal lymphatic collector [m-ELC] in C). A higher magnification micrograph (Figure 1D ①) shows the particularly prominent lymphatic vasculature of the efferent duct and its interconnection with that of the epididymal *caput*. It also points out the departure of the u-ELC to the PP surrounding the upper epididymal artery (u-EA). It can be noted that the lymphatic collectors coincide well with the segmentation of the organ. A *corpus* lymphatic collector (*Co*-LC) follows the body of the epididymis on the side adjacent to the testis connecting the *caput* with the *cauda* (Figure 1E). The inset within Figure 1E presents a higher magnification of the *corpus* lymphatic collector connections. This collector also connects to the main testicular lymphatic collector (TLC) at the *corpus* level (Figure 1E ②) as well as at the *cauda* level (Figure 1F ③). Also noticeable in the enlargements provided in Figure 1E and 1F is the presence of fluorescence under the capsule, faint in the *caput* (Figure 1E) but clearly more visible in the most distal segment (S10) of the *cauda* epididymis (Figure 1F and 1G). Finally, Figure 1G presents a global complex organization of the *cauda* lymphatic collectors. Overall, our observations revealed a very dense lymphatic network in the mouse epididymis.

### Complexity of the blood and lymphatic vascularization of the epididymis

Because fenestrated blood vessels also express VEGFR3, a marker specific for these vessels (PLVAP/Meca32) was used along with two additional markers for lymphatics, specifically LYVE1 and PDPN. As the 3DISCO organ clearing procedures resulted in a loss of the endogenous transgene (VEGFR3:YFP) fluorescence, anti-GFP was used to strengthen the signal. The concordance of the labeling obtained with the anti-GFP and an anti-VEGFR3 antibodies was controlled (see Supplementary Figure 1A). Similarly, Supplementary Figure 1B and 1C show positive controls of the multiplex labeling as well as controls with the secondary antibody, respectively.

Figure 2A and Supplementary Videos 1–3 show the immunoreactive vasculature after 3DISCO clearing of an adult mouse epididymis obtained after light-sheet imaging and IMARIS 3D reconstruction. The *caput* and *cauda* territories show intense PLVAP reactivity revealing a dense blood network. The *corpus* of the epididymis is also well vascularized, as can be seen in Supplementary Figure 2A and Supplementary Video 2. The video clearly shows that the lymphatic network is asymmetrical as it is very dense on the epididymal side next to the testis. In agreement with previous reports (Abe et al., 1984; Suzuki, 1982), the initial segment (IS) of the *caput* shows the highest reactivity (see Figure 2A and 2B as well as Supplementary Figure 2A and 2B), with each IS tubule strongly reactive at their periphery (Supplementary Figure 3A and 3B). Outside the IS, PLVAP reactivity is mainly found in the intertubular space as well as along the septum separating each epididymal segment. This is particularly true for the IS/S2–S3 and the S9/S10 septum junctions.

**Figure 2:**
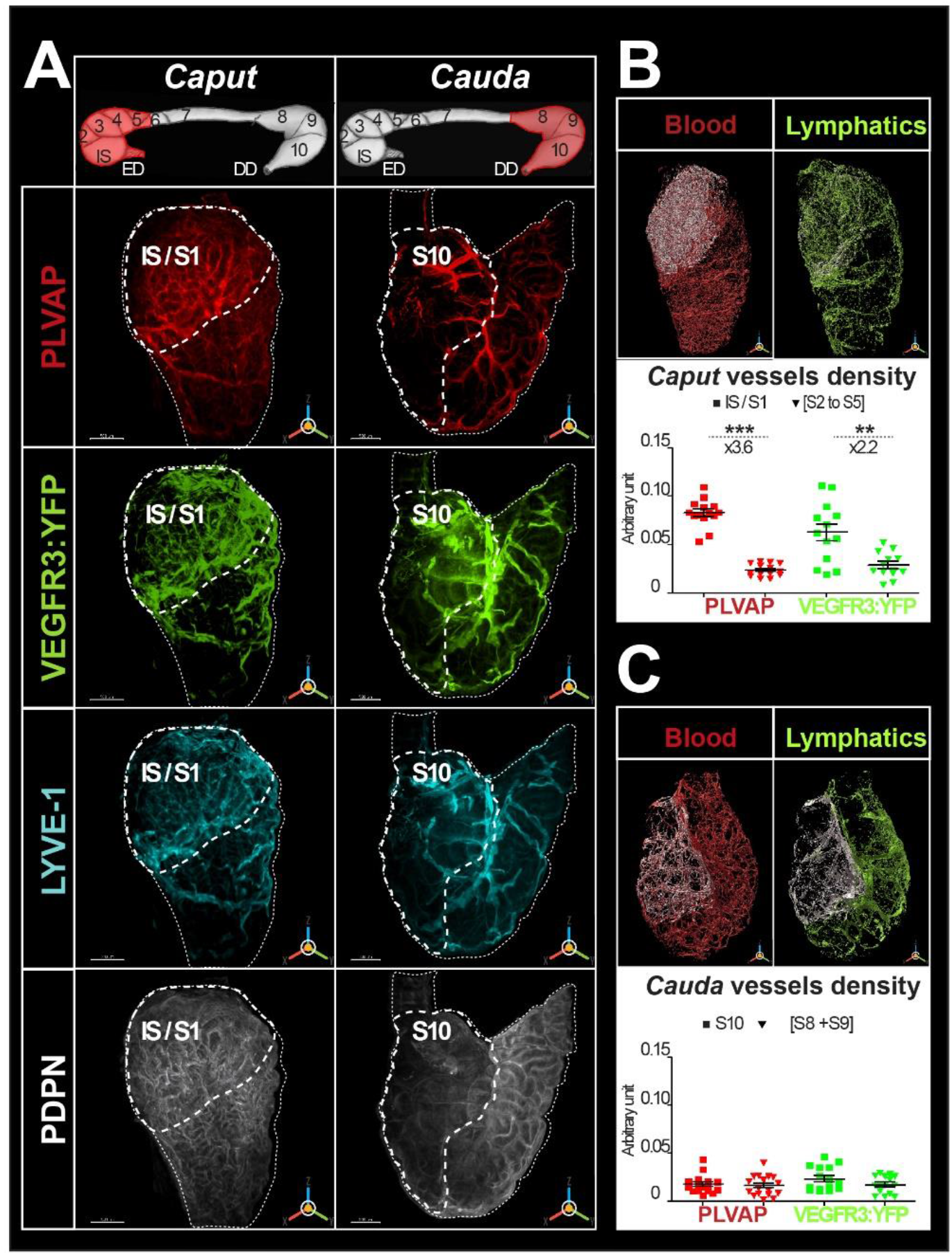
High-resolution three-dimensional (3D) imaging of the blood and lymphatic vasculature of the mouse epididymis after organ clearing. (A) The upper panels show a schematic representation of the mouse epididymis segmentation in which the *caput* (left) or *cauda* (right) regions have been highlighted (red). Representative multiplex immunostaining of the epididymis using four markers recognizing blood and lymphatic vessels is shown below each schematic. An anti-MECA32 antibody (red) reveals PLVAP^pos^ blood vessels, particularly fenestrated vessels. An anti-GFP antibody revealing the VEGFR3 :YFP transgene (green) as well as anti-LYVE1 (cyan) and anti-PDPN (white) antibodies were used as other lymphatic markers. After organ clearing, high-resolution 3D imaging was performed with a light-sheet ultramicroscope (LaVision BioTec). The two extreme segments, IS (initial segment or S1) and S10, are outlined by a large dashed line, while the *caput* (left) and *cauda* (right) regions are delineated by a small dashed line. The scale bar is 500 μm. Panel B shows the surface rendering of blood vessels (red, left) and lymphatic vessels (green, right) in the *caput*. The surface rendering of vessels in SI is in white in both cases. Vessel densities shown in the graphs below correspond to the ratio of the volume occupied by vessels in the SI or *caput* (minus SI) normalized to the total volume of the SI or *caput* (minus SI), respectively. Panel C shows the surface rendering of blood (red, left) and lymphatic (green, right) vessels in the *cauda*.Blood or lymphatic vessel densities arc measured as described in B for the *caput*. The Mann-Whitney test was used to determine statistical significance (*** p <0.0001, ** p<0.001, *p < 0.01, NS = not significant).

With the three lymphatic markers (VEGFR3:YFP, LYVE1, and PDPN), there is an intense and complex network (Figure 2A). This appears very similar in its organization to that of the blood vessels, especially in the IS (Figure 2A). However, outside the IS, lymphatic reactivity appears more extensive than that of the blood vessel marker (see Supplementary Figure 3B). As was the case for the blood vessel marker and as expected because fenestrated vessels express VEGFR3 (Partanen et al., 2000), lymphatic marker reactivity is stronger in the IS (Figure 2A). As seen in Figure 1, large external lymphatic collectors follow the septa, particularly at the IS/S2-S3 and S9/S10 boundaries. These collectors can be seen to dip inside the organ and irrigate the intertubular compartments, especially in the *cauda* region (see Supplementary video 3). The LYVE1 and VEGFR3:YFP profiles are very similar for the *caput* and *cauda* (Figure 2A and 2B). However, when viewed at higher magnification (Supplementary Figure 3A), they do not merge totally in either the *caput* or the *cauda* region, especially for the small lymphatic vessels separating the epididymal segments. The PDPN profile appears different from those of LYVE1 and VEGFR3:YFP (Figure 2A). The PDPN reactivity is more homogeneous, excludes the larger collectors, and is concentrated at the peritubular level (Figure 2A and Supplementary Figure 3A).

Figure 3A and Supplementary Figure 3A also show that the lymphatic marker (PROX1) gives very similar results to LYVE1 in the *caput* and *cauda* regions. However, the pattern of PROX1 is slightly distinct from that of VEGFR3:YFP and very different from that of PDPN. Interestingly, there are lymphatics expressing all four markers (VEGFR3:YFP, LYVE1, PROX1, and PDPN), particularly at the septa (Figure 3D).

**Figure 3:**
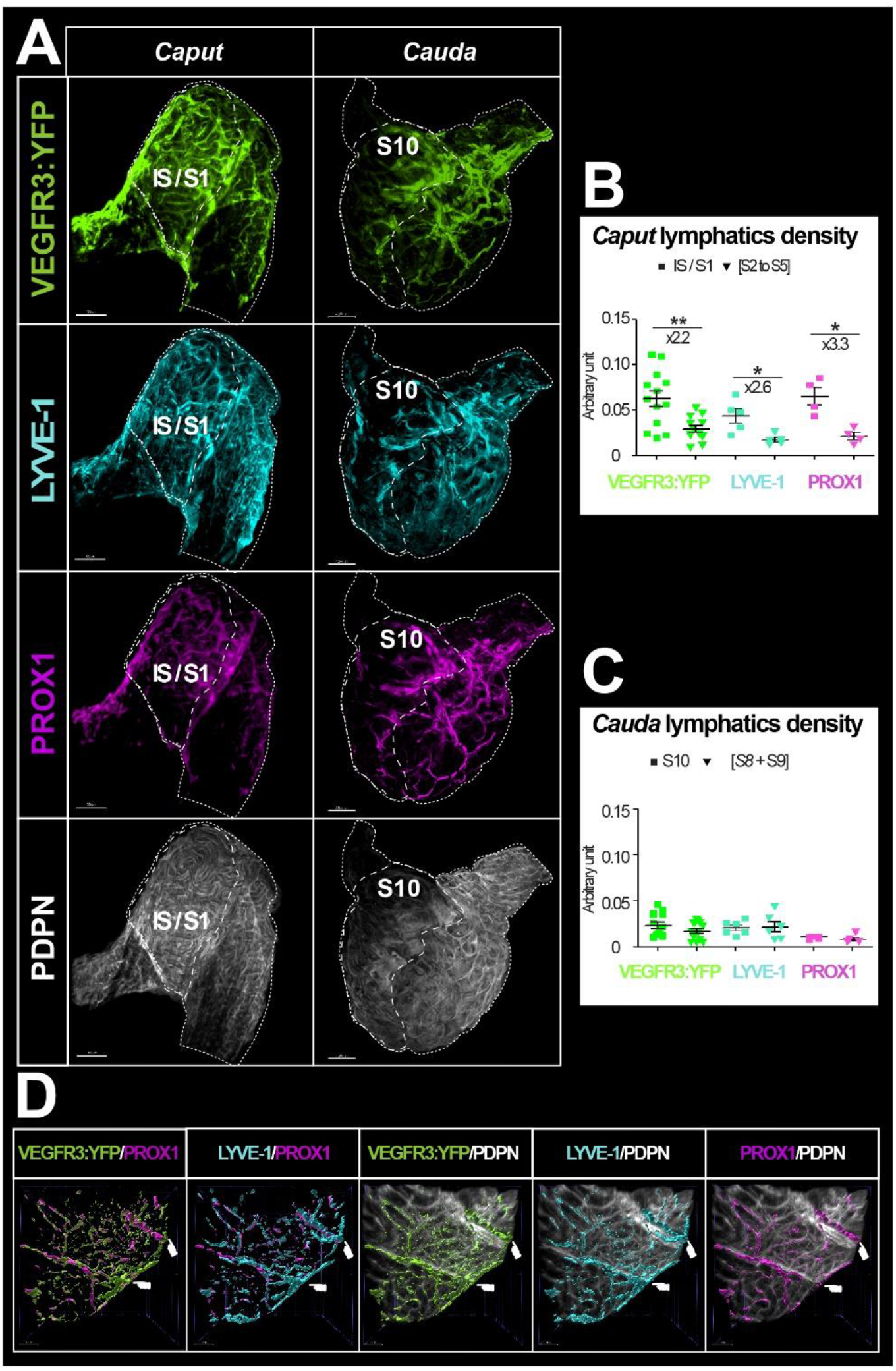
High-resolution three-dimensional (3D) imaging of lymphatics of the mouse epididymis after clearing revealed by multiplex immunostaining. Panel A shows whole-mount simultaneous immunostaining of the epididymis with four lymphatic markers: an anti-GFP antibody revealing the VEGFR3:YFP transgene (green), and anti-LYVEI (cyan), anti-PROXl (magenta), and anti-PDPN (white) antibodies. High-resolution 3D imaging was performed with a light-sheet ultramicroscope (La Vision BioTec). The two extreme segments, IS (initial segment of SI) and S10, are outlined by the large dashed line, whereas the caput (left) and cauda (right) regions are delineated by a small dashed line. The scale bar is 500 μm for the *caput* and 400 μm for the *cauda*. (B) *Caput* lymphatic density was determined for the different markers (as in Figure 2B), except for PDPN because of the nature of its signal. Panel C shows the same quantification obtained for tSIO and *cauda* (minus S10 region). The Mann-Whitney test was used to detennine statistical significance (*** p < 0.0001, ** p < 0.001, * p < 0.01, NS =not significant). Panel D shows a 3D region of interest of the *caput* region. Surface rendering of the lymphatics was performed as described previously using IMARIS software. The septa are indicated by the hand icon. The scale bar is 500 μm for the caput and 400 μm for die cauda.

We performed a surface assessment to better evaluate the density of the blood and lymphatic networks in the most proximal and distal epididymal segments (IS and S10) compared with that observed in the rest of the *caput* or *cauda*. To measure only the specific density of the lymphatic network, voxels with signal in the 555 nm channel (PLVAP/MECA32) were set to 0 in all other channels. The densities presented in Figures 2B, 2C, 3B, and 3C are the results of the vessel volume normalized by the total volume of the region of interest (IS, *caput*, S10, or *cauda*). As expected, PLVAP/MECA32^*pos*^ blood vessel density in the IS was approximately 3.5 times greater than in the rest of the *caput* (p <0.001). Similarly, the VEGFR3^pos^ lymphatic density was twice that of the rest of the organ (Figure 2B). A result of the same magnitude was obtained for LYVE1 and PROX1 (Figure 3B). Because of the expression profile of PDPN, surface rendering could not be used to calculate its density. In the *cauda*, there was no significant difference in the density of blood and lymphatic vessels. Comparison of vascular density between the *caput* (IS excluded) and the *cauda* showed a slight decrease in the *cauda* for both blood and lymphatic vessels (Supplementary Figure 2B).

### Dynamic development of epididymal vasculature related to organ maturation

To better understand the postnatal development of both networks, we followed their emergence during postnatal epididymal development from 10 to 40 DPN, when the organ is functionally mature (n = 5).

We present in Figure 4A (*caput*) and Supplementary Figure 4A (*cauda*) representative high-resolution images of the blood and lymphatic systems using three of the five markers presented above (PLVAP/MECA32, VEGFR3:YFP/GFP, and LYVE1) at 10, 20, 30, and 40 DPN. We used IMARIS software to estimate the IS volume relative to the *caput* volume (Figure 4B) and the S10 volume relative to the *cauda* volume (Supplementary Figure 4B) at each time point.

**Figure 4:**
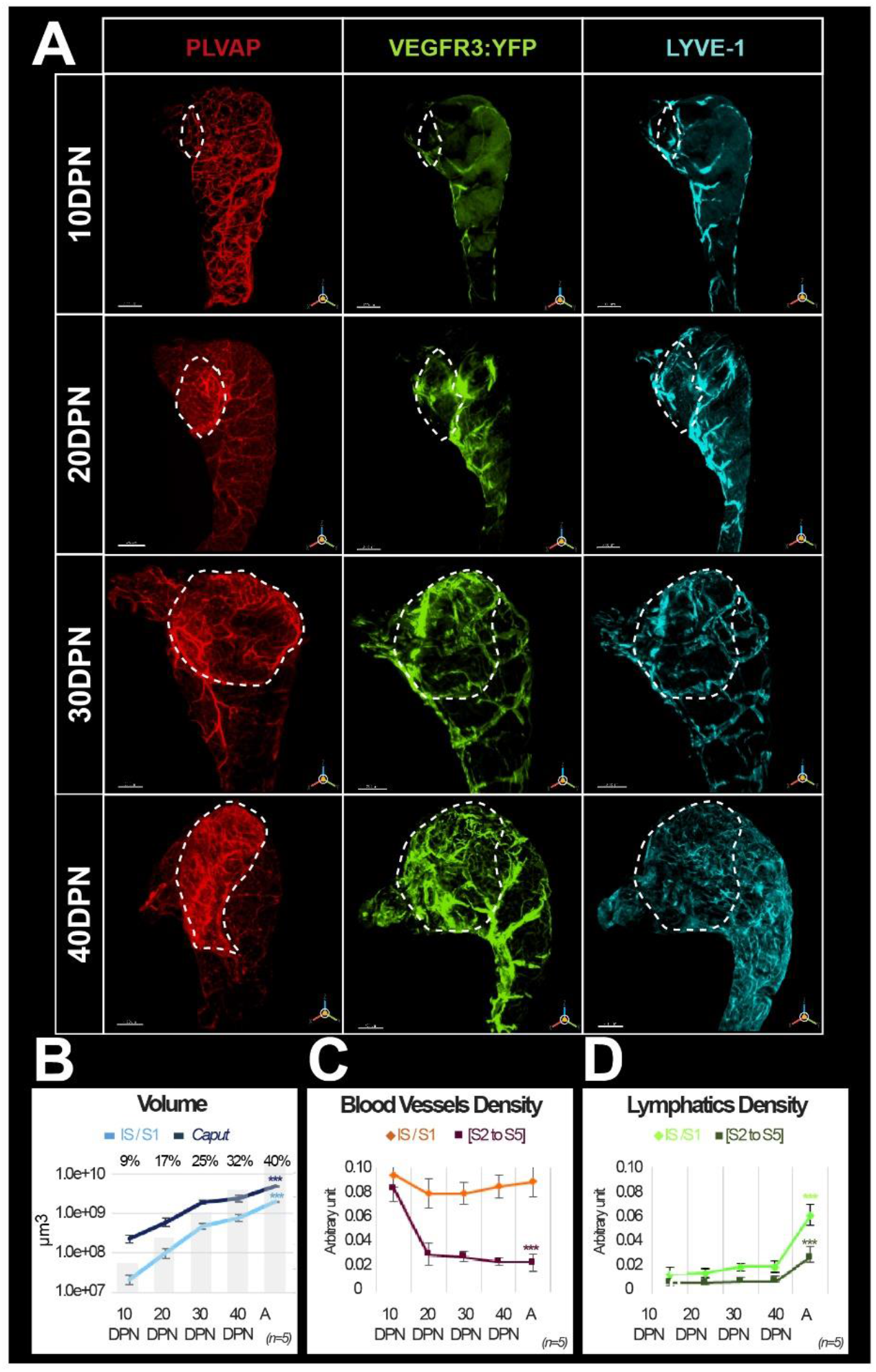
Evolution of the blood and lymphatic vasculature during postnatal epididymal development. Panel A shows representative three-dimensional (3D) images of blood and lymphatic networks at different postnatal stages during *caput* ontogenesis. Immunostaining was done with the blood vessel marker MECA32/PLVAP antibody (red) and the lymphatic marker LYVE1 antibody (cyan). The VEGFR3:YFP transgcnc is revealed by an anti-GFP antibody. The dotted line indicates the contours of the initial segment (IS = SI). DPN: days postnatal. The scale bar is 200 μm for 10 DPN, 300 μm for 2 0 and 30 DPN, and 400 μm for 40 DPN. Panel B shows the evolution of the volume (in log10) of the SI segment (light blue curve) and the *caput* region (dark blue) during postnatal development. The superimposed histogram gives the proportion of volume occupied by the SI relative to the *caput* region at different stages of postnatal development. Surface rendering of blood vessels and lymphatics was performed using IMARIS software. Blood and lymphatic vessel densities were calculated as described previously (Figures 2 and 3). Quantification was performed on five replicates for the different postnatal developmental stages. Panels C and D arc graphs presenting densities (mean and standard error of the mean) of blood and lymphatic vasculature, respectively. The Kruskal-Wallis test with Dunn’s posttest correction was used to determine statistical significance (*** p < 0.0001, **p < 0.001, *p < 0.01).

IS progresses 4 times faster than the *caput* between 10 DPN and adulthood (A), representing less than 10% of the *caput* at 10 DPN whereas it represents 40% of the adult *caput* (Figure 4A and 4B). Following the stronger postnatal development of the IS, we observed that the *caput* vascularization is rather homogeneous at 10 DPN, whereas from 20 DPN onwards, the blood vessels seem to concentrate in the IS segment. This is also well illustrated in the densitometric quantification shown in Figure 4C. This figure also shows that the density of PLVAP^*pos*^ blood vessels in the IS segment is rather stable during postnatal development despite the increase in the segment volume. In contrast, there is a strong decrease between 10 and 20 DPN in the rest of the *caput*. In the *cauda* region (see Supplementary Figure 4A–C), PLVAP^*pos*^ blood vessel expansion follows organ growth without a specific segmental pattern. However, as was the case in the *caput*, between 10 and 20 DPN, there is a decrease in PLVAP^*pos*^ blood vessel density in the caudal region.

Looking at the two lymphatic markers (VEGFR3:YFP and LYVE1), we observed very similar kinetics of postnatal lymphatic development in the *caput* (Figure 4A) and *cauda* (Supplementary Figure 4A). There is a progressive increase in VEGFR3:YFP/LYVE1 lymphatic vessels that follows postnatal organ growth in a fairly linear fashion at least between 10 and 40 DPN (Figure 4D and Supplementary Figure 4D). Beyond 40 DPN, there appears to be a greater expansion of lymphatics in the *caput* and *cauda*, in the IS and S10, respectively, compared with the rest of the organ (Figure 4D and Supplementary Figure 4D). This is particularly true in the IS because the size of this territory is increased approximately 4 times between the 10 and 40 DPN stages.

Because blood and lymphatic vascular density data suggest neovascularization during postnatal epididymal development, we performed localization and quantification of angiogenic (VEGF-A) and lymphangiogenic (VEGF-C and VEGF-D) factors, whose receptors are VEGFR2 (also known as KDR and FLK1) and VEGFR3, respectively (Figures 5 and 6). Quantification of the VEGFR ligands (VEGF-A, VEGF-C, and VEGF-D) was performed by western blotting as shown in Figure 5A. For the angiogenic ligand (VEGF-A), there was a decrease between 10 and 20 DPN followed by a plateau, and then an increase at 40 DPN; by adulthood, VEGF-A had returned to the initial level (Figure 5B). For the lymphangiogenic ligands (VEGF-C and VEGF-D), we observed a linear increase from 30 DPN for both ligands, consistent with the accumulation of their corresponding receptor VEGFR3 (Figure 5B). An analysis of covariance of the different ligand isoforms of VEGF (see Supplementary Figure 5) with respect to the VEGFR3 receptor (see Figure 5C) shows strong statistical significance for the active ligand isoforms (i.e., the 21 kDa form for VEGF-C and the 24 kDa form for VEGF-D). In addition, the tissue localization of VEGFR2 is very similar to that of VEGFR3 (Figure 6A), being present mainly in a peritubular position at the level of the IS whereas in the more distal part of the organ it seems to be restricted to some interstitial vessels (used with permission from Dr. A. Medvinsky; VEGFR2:GFP mouse strain; Xu et al., 2010). In accordance with previous work (Korpelainen et al., 1998), the tissue localization of its angiogenic ligand (VEGF-A) seems to correspond correctly (Figure 6A), supporting the intense blood vascularization observed in this very proximal segment of the epididymis. Both of the lymphangiogenic ligands—VEGF-C and VEGF-D—show a similar tissue localization, being weakly present in the IS and increasingly present when moving toward the tail of the epididymis, mainly in an interstitial position (Figure 6B). A co-localization (Pearson coefficient = 0.684 ± 0.038) and co-occurrence analysis (Mander coefficient VEGF-C = 0.934 ± 0.028 and VEGF-D = 0.569 ± 0.031) supports the hypothesis of heterodimerization of the two ligands.

**Figure 5:**
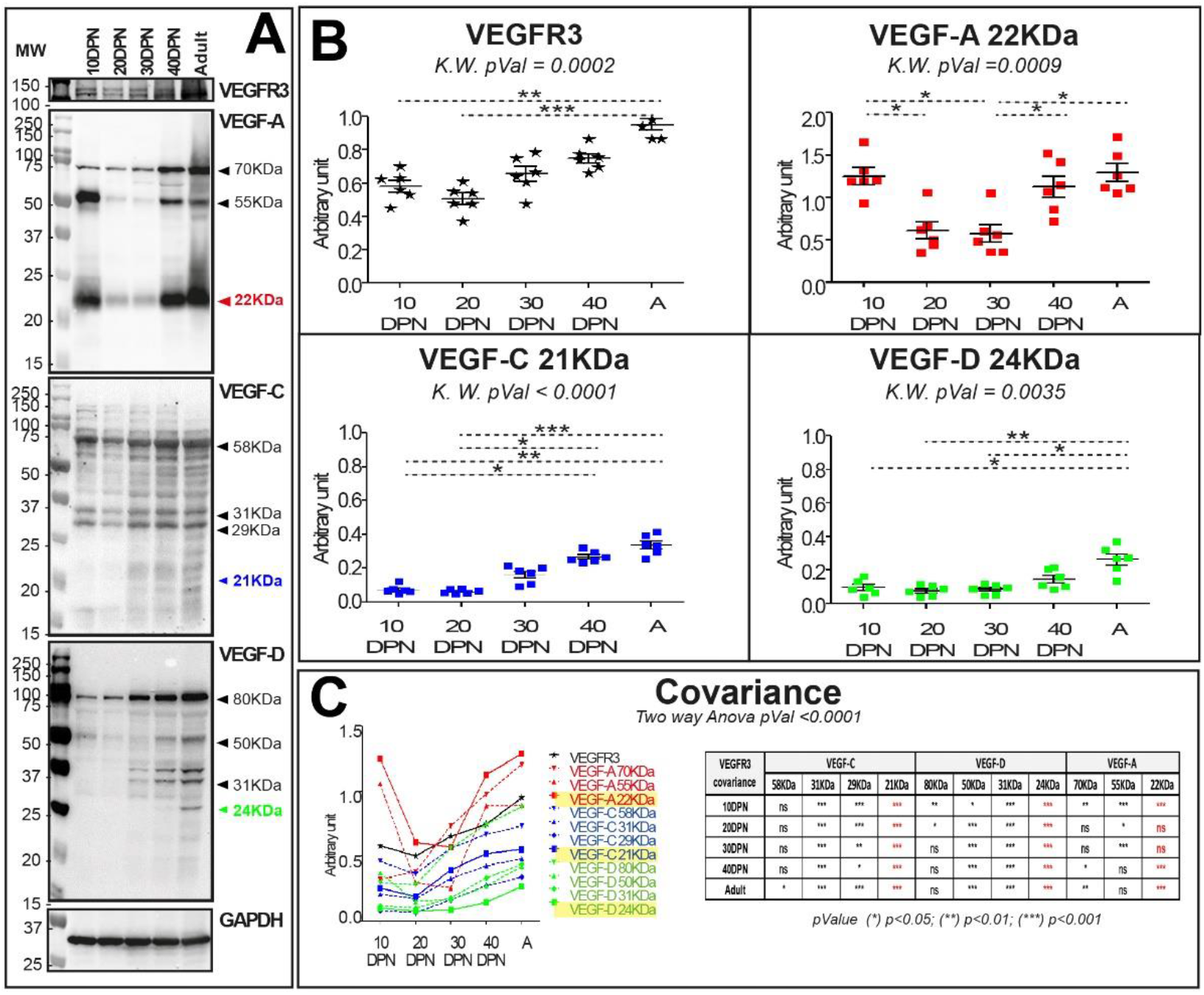
VEGF-A, VEGF-C, and VEGF-D levels vary during postnatal epididymal development. Panel A shows the expression profile obtained in total extracts of epididymal proteins at different stages of development. The profiles obtained arc presented in the following order: VEGFR3, VEGF-A, VEGF-C, and VEGF-D. GAPDH was used for normalization. Panel B shows the quantification of epididymal proteins extracted from six mice. Quantification of VEGFR3 is shown in black, and VEGF-A, VEGF-C, and VEGF-D are shown in red, blue, and green, respectively. The Kruskal-Wallis test and Dunn’s posttest correction were used to determine statistical significance (*** p < 0.0001, **p < 0.001, *P < 0.01). Panel C shows the comparison of different isoform profiles. Covariance is assessed by a two-way analysis of variance.

**Figure 6:**
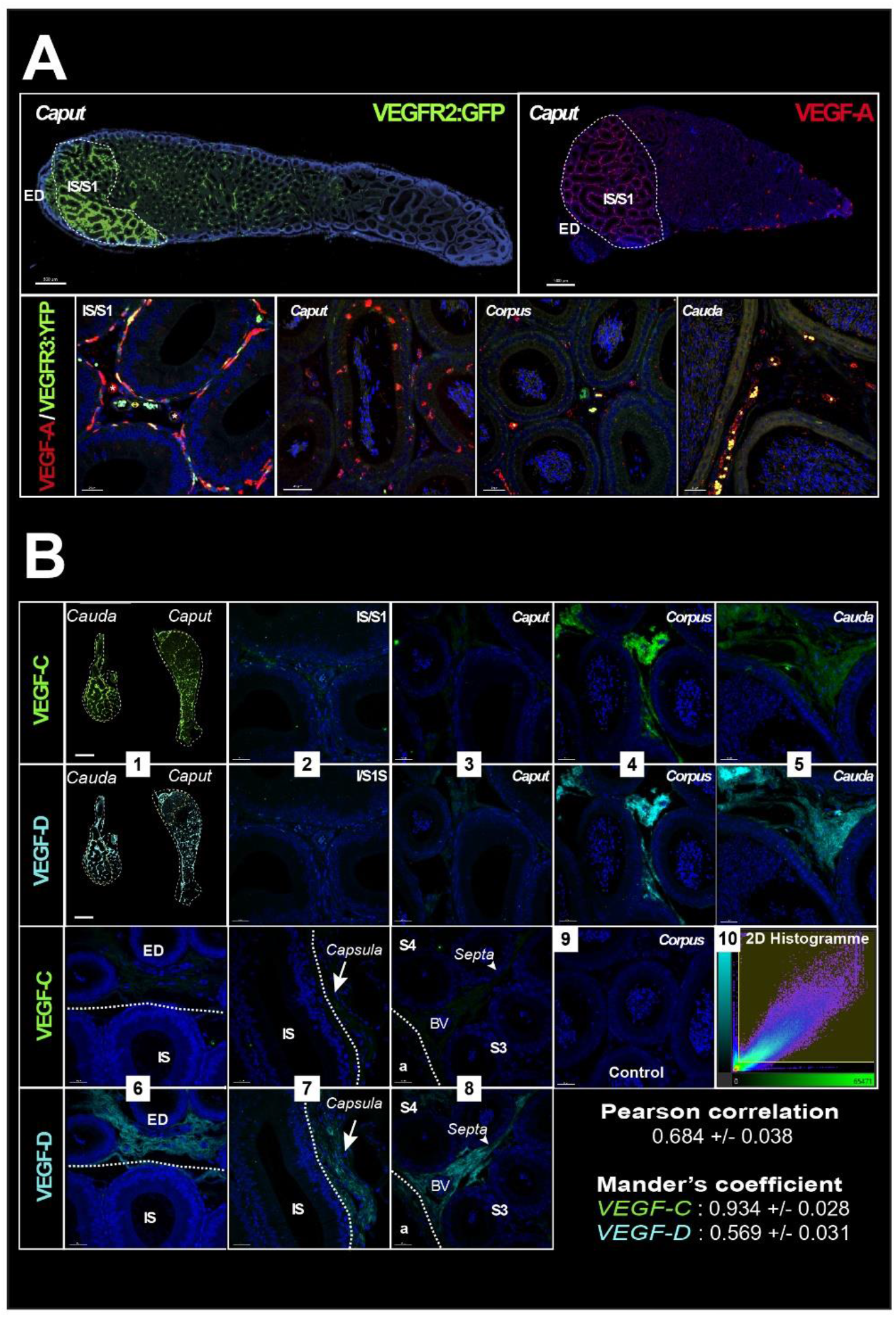
VEGF receptors and ligands in the mouse epididymis. Panel A (upper row) shows images (Zeiss Axio-Imager) of the expression of the VEGFR2:GFP transgene (green) and its ligand VEGF-A (red) (used with permission from Prof. A. Medvinsky) obtained from paraffin sections of adult epididymis. The lower row shows confocal views (SP8, Leica) of the same VEGF-A labeling. Panel B shows in (1) a mosaic view (AxioVison scanner) of an epididymis after immunolabcling with VEGF-C (green) and VEGF-D (cyan). Photographs 2-9 were taken with a confocal (SP8, Leica) at the levelof the initial segment (IS)/Sl (2), *caput* (3), *corpus* (4), and *cauda* (5). Notable differences in expression of these two ligands are shown in photographs 6-8, which respectively concerns the efferent duct and the 1S/S1 boundary, the capsule, and a septum. Photograph 9 is a negative control. Pearson correlation and Mandcr’s co-occurrcncc were used to analyze the relation between the two ligands withthc IMARIS co-localization module. The values shown represent means with standard error of the mean.

To conclude, we show in Figure 7A (upper panels) a dynamic 3D view of the *cauda* epididymis lymphangiogenesis as well as our interpretation of what is happening, presented in the lower panels. The schematic, shown in Figure 7B, presents the detailed organization of the epididymis lymphatics with respect to the tubules, septa, and orientation of the organ, with the anterior side of the epididymis tubule in close contact with the testis. LYVE1^pos^ cubblestone cells are observed at 10DPN while very few VEGFR3-YFP^pos^ cells are visible. Later, cubblestone cells appear to aggregate to form tubular structures. Concomitantly, we record an increase in VEGFR3-YFP^pos^ structures associated with a decrease in PLVAP^pos^ vessels. This kinetics of events is consistent with active lympho-vasculogenesis during postnatal epididymal development.

**Figure 7:**
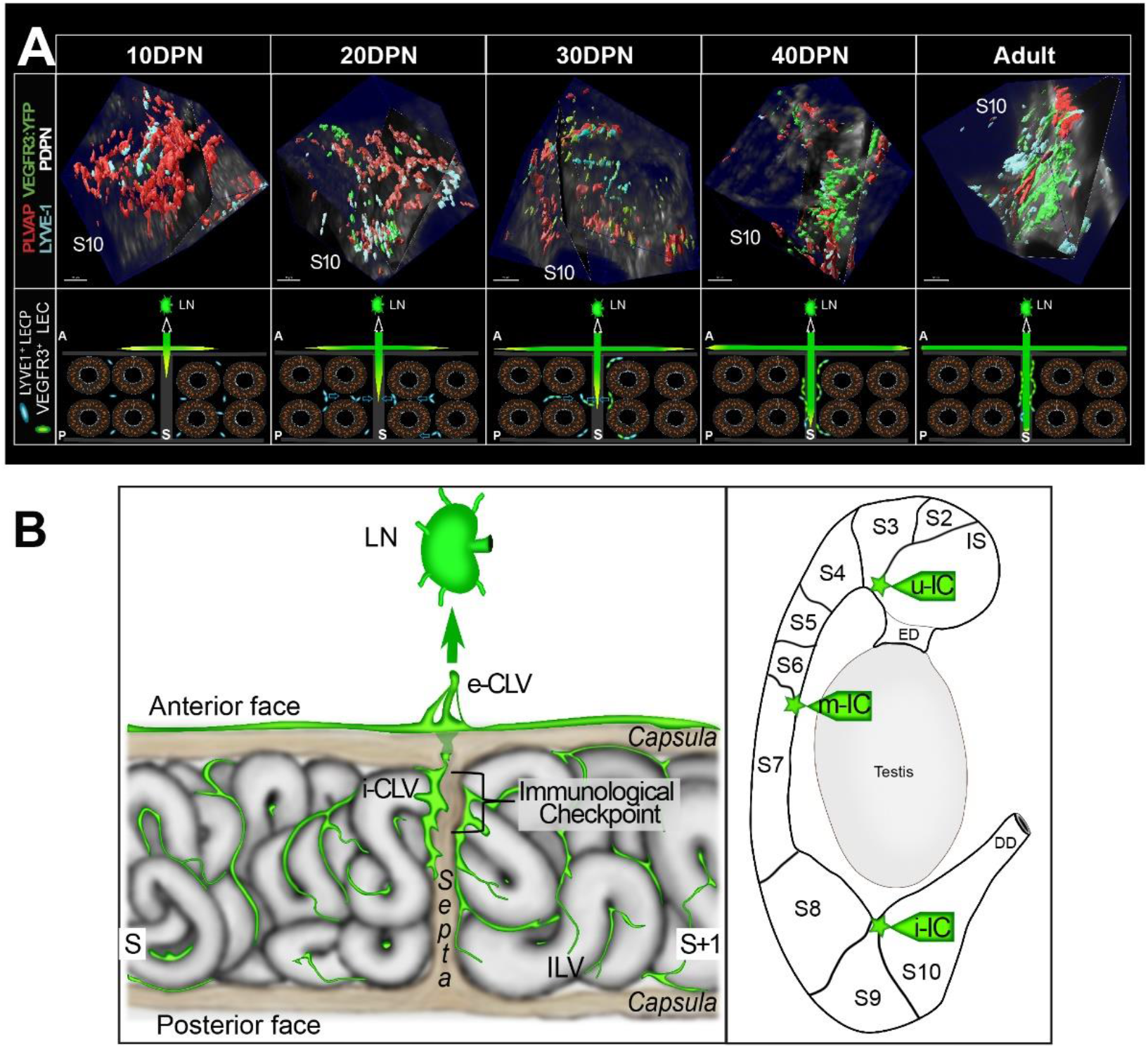
Dynamic evolution of the external and internal blood and lymphatic vasculature during postnatal epididymal development. Panel A (upper row) shows a representative three-dimensional (3D) view of the *cauda* region at the S8-S9/S10 septum, during postnatal development of the epididymis, after surface rendering. Images were obtained using the light-sheet ultramicroscope with a 20× objective lens. Putative LYVE1 ^+^ lymphatic precursors are in cyan and VEGFR3:YFP lymphatics are in green, while PLVAP^+^ blood vessels are in red. The black plane that divides the image is positioned at the septum level. PDPN expression is in white and helps visualize the folded tubule. The lower row shows our interpretation of the events observed concomitantly with a progression via lymphangiogcncsis of peripheral lymphatics that progress and radiate into the organ at the level of the septum, from the anterior side (adjacent to the testis) toward the posterior side of the epididymis. Lymphangio-vasculogenesis is represented by interstitial LYVE1^+^ lymphatic endothelial cell progenitors (LECP) (cyan) that aggregate and migrate to join VEGFR:YFP^+^ lymphatic endothelial cells (LEC), which ultimately organize into a lymphatic (green). For the sake of clarity, PLVAP^+^ blood vessels have not been represented here. (A =anterior side adjacent to the testis, P =posterior side, S =septum, LN =lymph node). Panel B presents our view of the lymphatic vasculature at the level of a “checkpoint” septum. The large collectors enter at the septum into the epididymis and radiate within the adjacent segments at the interstitial level, where initial lymphatics are found. The septum where the tubule crosses from one segment to another one would be the most “monitored” site and therefore the richest in lymphatics. e-CLV = external collector lymphatic vessel; i-CLV = internal collector lymphatic vessel; ILV = initial lymphatic vessel; LN = lymph node; S = segment.

## Discussion

To the best of our knowledge, the data presented in this study represent the first in-depth simultaneous observation of blood and lymphatic networks in the mammalian epididymis, both in the adult stage and during postnatal epididymal ontogeny. With the combination of lymphatic markers used in this work, we have shown that a dense network of conventional lymphatics can be observed in the epididymis, with initial lymphatics present in the interstitial compartment and collecting lymphatics present at the septa. Reports describing mammalian epididymal lymphatics are rare and rather old (Perez-Clavier et al., 1982; McDonald, 1988). The technology used was not very efficient and consisted mainly of ink or contrast agent injection. It concerned only the peripheral epididymal lymphatic network with the organization of the collectors connecting the PP. More recently, Hirai and colleagues (2010) performed the first histological study of mouse epididymal blood and lymphatic vasculature using the markers CD31/PECAM1 and LYVE1, respectively. Although PECAM1 is used to visualize blood vessels, it is not an exclusive blood marker because it is also found expressed at button-like junctions of endothelial cells in the initial lymphatics (Privratsky and Newman, 2014). Furthermore, to characterize lymphatics, the use of LYVE1 alone limits the power of the analysis because this marker recognizes only a subpopulation of all lymphatics (Kato et al., 2006). In addition, conventional two-dimensional (2D) light microscopy combined with a horseradish peroxidase (HRP) reporter on paraffin-embedded tissue sections did not allow for a high level of accuracy in providing a holistic view of a structured network. Therefore, although pioneering, Hirai et al.’s (2010) study was limited and lacked the power that we have introduced by using whole-mount multiplex immunolabeling, organ clearing, ultramicroscopy imaging, and 3D reconstruction with a highly adapted lymphatic-YFP transgenic reporter mouse model. The sensitivity and power of our study has allowed us to present for the first time an in-depth analysis of blood and lymphatic networks both in the adult and during postnatal epididymal development.

We have shown that the lymphatic network is denser on the anterior face of the epididymis (face adjacent to the testis) and progresses in the organ toward the posterior face. We found that the collecting lymphatics are mainly observed on the anterior surface and radiate inwards toward the posterior surface following the septa. The initial lymphatics drain the interstitial compartment and join the collectors at the septa, resulting in an asymmetric distribution (anterior/posterior) of lymphatic vascularization. This centrifuge development of the lymphatic network is easily visible (Figure 4A and Supplementary Video 2). This tree structure is logical if we consider that lymphatic vascularization progresses during mouse development following the PP before reaching the *rete testis* and sprouting laterally, as had been suggested earlier (Svingen et al., 2012). Each epididymal septum has lymphatic drainage, but three septa appear to be more involved, including the S1/S2–S3, S6/S7, and S8–9/S10 junctions, with the latter having the largest lymphatic collector. We expected this finding because it corresponds to the position of the arteries and veins that irrigate the epididymis (Suzuki, 1982), and it is known that the large lymphatic collectors follow the path of the arteries (Gruffaz, 1984). Previous researchers have hypothesized that this strict epididymal segmentation is involved in the control of ascending pathogens, although they have not provided clues as to how this might be accomplished (Stammler et al., 2015; Turner et al., 2003). Supporting this hypothesis is the observation that large lymphatic collectors that are in direct and rapid connection with LNs and the blood supply that allows for the arrival of leukocytes are present at these sites. These three septa could then be considered important checkpoints. This view corroborates recent data showing that immune responses in autoimmune epididymitis are concentrated in the epididymal *cauda* and *corpus*, where we do see the major lymphatic collectors (Wijayarathna et al., 2020). Looking at the postnatal development of the epididymal lymphatic network, our data show a progressive increase that follows the postnatal growth of the organ between 10 and 40 DPN. Beyond this time point, the lymphatic network progresses more in the S1 and S10 compartments. This argues for active lymphangiogenesis or/and lymphangio-vasculogenesis, the two modalities by which lymphatics develop, in the *caput* and *cauda* territories. Lymphangiogenesis corresponds to the extension of existing vessels, also known as “sprouting,” while lymphangio-vasculogenesis involves the participation of precursor cells that create clusters later integrating already formed lymphatic vessels (Gutierrez-Miranda and Yaniv, 2020; Jafree et al., 2021). Lymphangio-vasculogenesis has been reported in the development of the mesenteric lymphatic vessels (Stanczuk et al., 2015) and for the heart and kidney lymphatic networks (Jafree et al., 2019; Lioux et al., 2020), two organs with a septal organization. The lymphatic neovascularization we observed during postnatal epididymal development is supported by the observation that lymphangiogenic ligands (VEGF-C and VEGF-D) and their corresponding receptor (VEGFR3) are concordantly present in the developing tissue. Active lymphangio-vasculogenesis is also supported by our observation that foci expressing LYVE1 can be detected between 10 and 20 DPN, and eventually extend and join the initial lymphatics (see Figure 7A).

In agreement with what has been reported previously (Suzuki, 1982), a testis-like blood network is present in most of the epididymis (segments 2–9) with intertubular vessels connected by perpendicular micro-vessels in a rope ladder organization. Distinctly, in the most proximal (IS/S1) and distal (S10) segments of the epididymis, we found micro-vessels encircling each tubule in a spider web organization with capillaries that penetrate the epithelial sublayer. Because the IS and S10 compartments of the epididymis have distinct embryonic origins and are functionally different (Abe et al., 1984; Johnston et al., 2007), it is difficult to explain this common vascular organization. We hypothesize that the common set-up of these two territories where capillaries penetrate the sub-epithelial layer is associated with their immune function. The S10 compartment must monitor ascending pathogens and maintain self-tolerance to sperm antigens that accumulate in this storage part of the organ. Similarly, the S1 compartment is the final gatekeeper preventing ascending pathogens from reaching the immune-privileged seminiferous tubules, while at the same time it must be tolerant to sperm-specific antigens entering the epididymal tubule and considered non-self. Therefore, in both compartments, a spider web blood network can be expected to allow immune cells easy access to the epididymal tubule. This should be particularly true for the S1 segment because, independently of infectious situations, it is constantly solicited in the adult animal by the arrival of sperm antigens and by its function of reabsorption of Sertolian fluid requiring very permeable vessels. This hypervascularization of the S1 epididymal compartment is further supported by the fact that it has been shown to be PLVAP^*pos*^, a marker of fenestrated vessels allowing high trans-endothelial transport (Guo et al., 2016). In other contexts, PLVAP has been shown to be required for fenestron biogenesis and organization, controlling fenestron permeability, angiogenesis, as well as leukocyte and antigen migration (Rantakari et al., 2015). In addition, the presence of VEGFR2, VEGFR3, and VEGF-A in peritubular blood vessels of the S1 compartment argues for active angiogenesis (Bussolati et al., 2003). This has been corroborated by the recent single-cell data (Rinaldi et al., 2020) showing that expression of angiogenic markers such as those mentioned above occurs in endothelial cells of the caput mouse epididymis. While active angiogenesis during postnatal epididymal development makes sense because it accompanies postnatal growth of the organ, it is more surprising that angiogenesis is still active in the adult IS epididymal compartment. In adult tissues, angiogenesis is most commonly associated with vascular remodeling in tissue repair processes and tumor progression (Komi et al., 2020). To date, self-renewal of the epididymal epithelium is collectively assumed to be rather slow, a situation that should not be associated with high angiogenic activity. However, very recent data could modify this general assumption because it has been reported that the basal cells of the epididymis share common properties with adult stem cells (Dufresne et al., 2022). This could support the idea that the epithelium of the adult epididymis may be in a perpetual process of self-renewal that could be promoted by the quasi-inflammatory situation inherent to immune stimulation mediated by sperm antigens.

It is interesting to note that during postnatal ontogeny of the epididymis, PLVAP expression increases in the S1 territory but decreases in the other regions of the organ as early as 15 DPN. Whether this is due to blood vessels loss or to PLVAP^*pos*^ permeable fenestrons becoming PLVAP^*neg*^, less permeable vessels will require further study. In the mouse model, it has been shown that the development of the S1 territory occurs between 15 and 20 DPN concomitantly with the arrival of testicular fluid (Jegou, 1982), which has been shown to stimulate the Raf/Mek/Erk pathway (Xu et al., 2016). Therefore, it is possible to consider that VEGF-A could be one of the lumicrine factors triggering angiogenesis and differentiation of S1/IS. In agreement with this, it has been reported that mice overexpressing VEGF-A showed dilation of the *caput* and *corpus* epididymis, with enlarged and more permeable blood vessels (Korpelainen et al., 1998).

More recently, Pawlak and Caron (2020) have suggested that such vessels could be hybrid lymphatics expressing both blood and lymphatic markers. Such hybrid vessels have been found in distinct vascular beds, including high endothelial venules (HEV) in secondary lymphoid organs, liver sinusoidal endothelial cells (LSEC), Schlemm’s canal endothelial cells (SCEC) of the eye, the ascending vasa recta (AVR) of the kidney inner medulla region, and the remodeled spiral arteries (RSA) of the placenta decidua (Gola et al., 2020; Kenig-Kozolvsky et al., 2018; Kim et al., 2017; Pawlak et al., 2019; Russell et al., 2019; Stamataki et al., 2020) where they perform specialized exchange functions. As shown in Supplementary Figure 7, these structures share many markers, including those tested in the present work, suggesting that some of the assumed blood vessels found in the S1 epididymal territory might be hybrid vessels. Given the close mesonephric origin and similar fluid reabsorption function of the IS/S1 epididymal territory and the AVR, it is possible that the two structures share such hybrid vessels. Tie2 immunohistochemistry (not shown) that has been shown to be a marker of all currently described hybrid lymphatics (Kenig-Kozlovsky et al., 2018), supports this hypothesis.

In conclusion, based on our extensive study of the blood and lymphatic vascularization of the epididymis in mice, we propose that the epididymis presents two types of lymphatic vessels. On the one hand, there are peritubular PLVAP^*pos*^ hybrid vessels, very permeable, particularly well represented in the S1 compartment that, according to our hypothesis, participate both in the reabsorption function and in the immune surveillance of this territory. On the other hand, we have more conventional lymphatics mainly in the interstitial compartment. The various segments of the epididymis are drained by initial and collecting lymphatic vessels, the latter being closely associated with the septa and then connecting via the PP to the most proximal LN. Three checkpoints are represented by the IS/S2–S3, S6/S7, and S8–9/S10 septa. It is not yet clear to us that the lymphatic drainage of the S8–9/S10 compartment goes to the PP because it could also be drained by the lymphatic circuit monitoring the vas deferens. This detailed knowledge of the blood and lymphatic circuits of the epididymis highlights how open the epididymis is to the systemic compartment and how different its immune control is from that of the testis, where lymphatic collectors are restricted to the conjunctiva and never reach the interstitial compartment (Hirai et al., 2012; Svingen et al., 2012). If these observations in the mouse model are somehow translatable to the human epididymis, they could help to understand the complex inflammatory and immune contexts of post-testicular sperm maturation and their impact on male fertility.

---

**Supplementary Figure 1:**
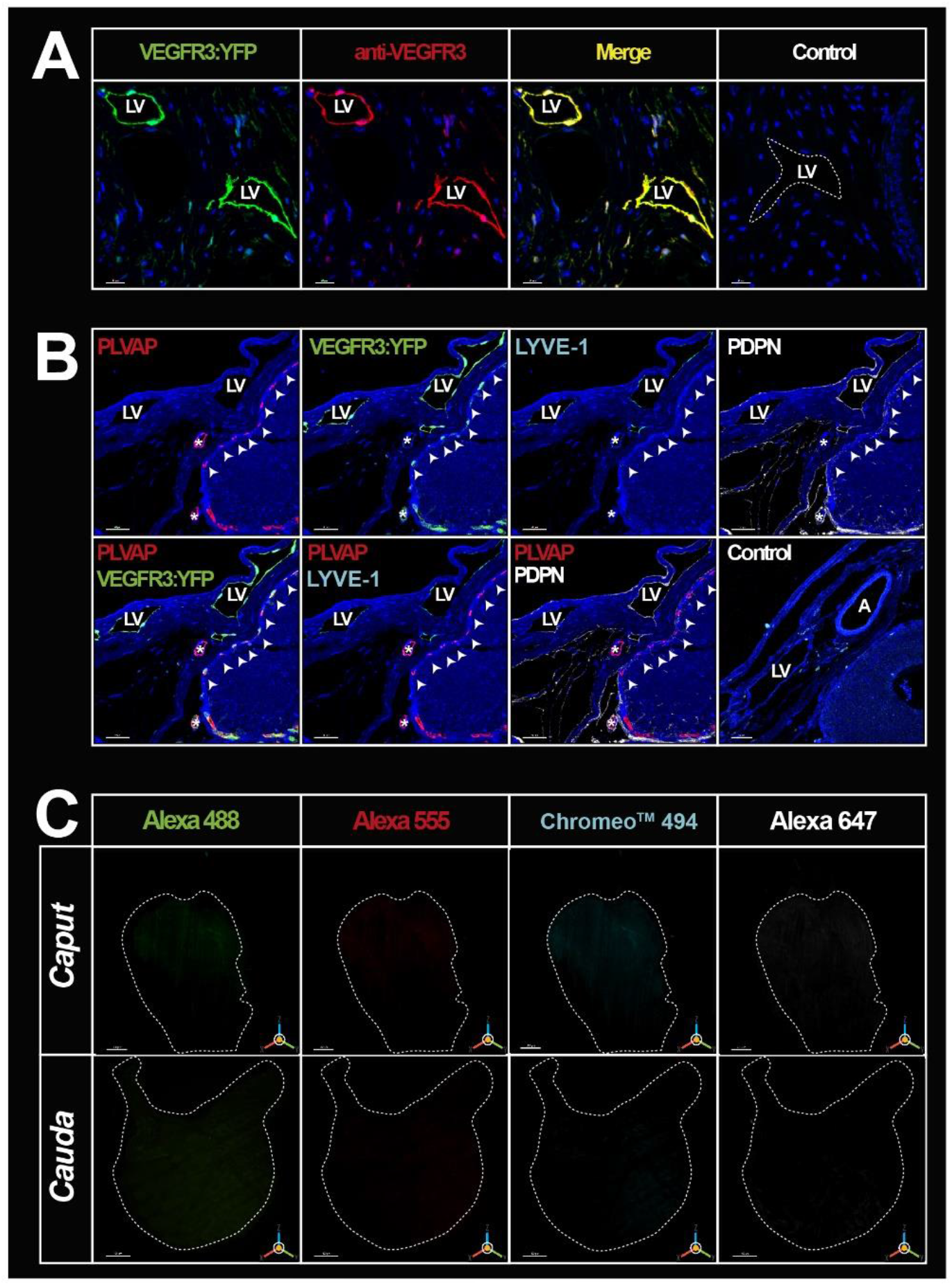
Positive and negative immunohistochemical controls. Panel A shows confocal views of multiplex labeling with the anti-GFP antibody (green) and the anti-mouse VEGFR3 antibody (red)on paraffin sections of an adult mouse epididymis, attesting that the two antibodies recognize the same structures. The scalebar is 20 μm. Panel B shows the compatibility of the four markers/dyes chosen for imaging blood and lymphatic vessels. Multiplex paraffin immunodetection of PLVAP/Alexa A555 (red), VEGFR3:YFP/GFP/Alcxa A488 (Green), LYVE1/Chromco™ 494 (cyan), and PDPN/Alexa A 647 (white) at the junction between the initial segment (IS)/Sl and the epididymal capsule. We noted the presence of fenestrated PLVAP^pos^ blood vessels at the periphery of the epididymal tubule of the IS (white arrowheads). These arealso positive for VEGFR3:YFP/GFP but negative for ?YVE1 and PDPN, demonstrating that there is no cross-detection between the labeling revealed with ChromcoTM 494 and those revealed by Alexa 488, Alexa 555, and Alexa 647. The peripheral lymphatic vessels of the organ located in the capsule (LV) are negative for PLVAP and positive for the three lymphatic markers (VEGFR3:YFP/GFP, LYVE1 and PDPN). We noted the presence of intertubular blood vessels (*), which present heterogeneous labeling. The upper one is only PLVAP^pos^ while the lower one is PLVAP^pos^ and VEGFR3 :YFP^pos^. In addition, note that the latter shows peripheral labeling for PDPN, while that is not the case for the upper one. A =Artery. Panel C shows three-dimensional (3D) imaging obtained with a light-sheet ultra-microscope (LaVision Biotech) of the clarified epididymis *caput* and *cauda* after whole-mount incubation with the different dyes used to reveal the blood and lymphatic markers (Alexa 488 [green], Alexa555 [red] ChromcoTM 494 [cyan], and Alexa 647 [white]).

**Supplementary Figure 2:**
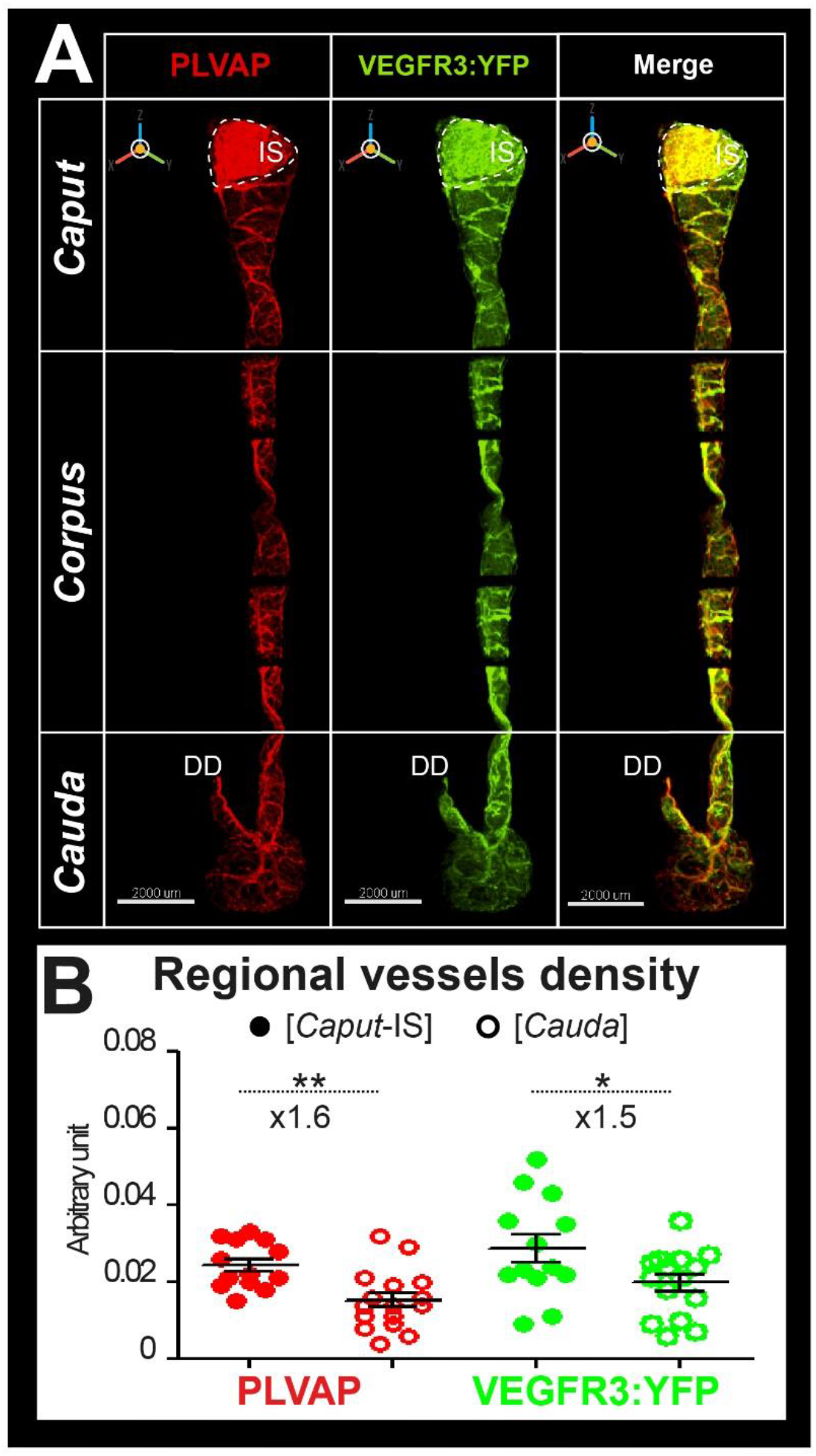
Three-dimensional (3D) view of blood and lymphatic vascularization of whole epididymis after 3D1SCO clarification. Panel A shows representative 3D views of an epididymis of an adult VEGFR3:YFP transgenic mouse obtained by light-sheet ultramicroscopy (LaVision Biotech, 2× resolution) after immunodetection using the blood marker Meca32/PLVAP (red, left panel) or the lymphatic VEGFR3:YFP transgene, revealed using an anti-GFP antibody (green, middle panel). The right panel presents a merged view. Panel B presents a comparison of blood and lymphatic vessel density between the caput and cauda as described in the Figure 2 legend. The Mann-Whitney test was used to determine statistical significance (** p < 0.001, * p <0.01).

**Supplementary Figure 3:**
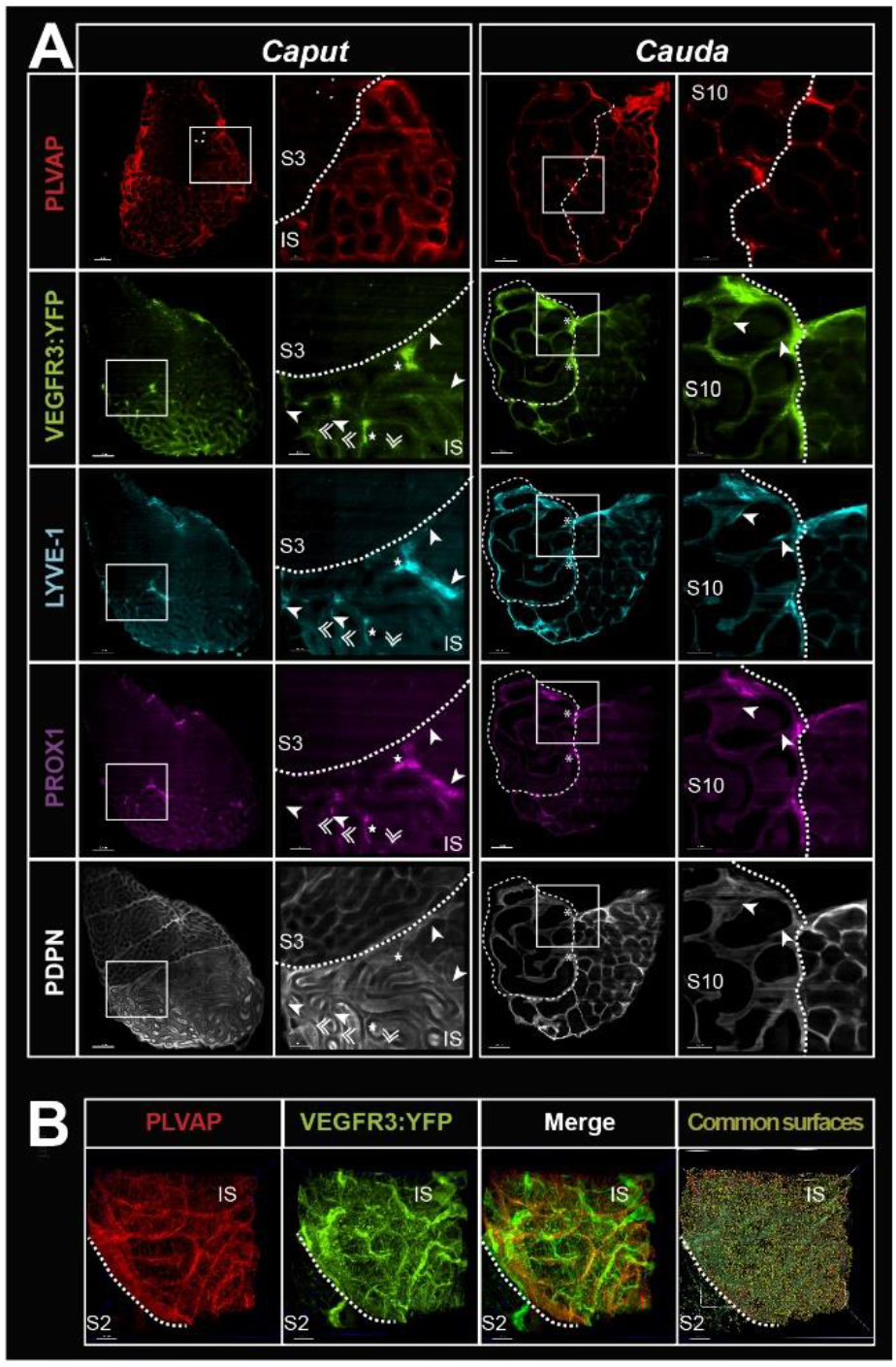
Cross-sectional views of the clarified caput and cauda epididymides. Panel A presents median slices of the caput (left) and cauda (right) shown in Figure 2 for the PLVAP blood marker (red) and in Figure 3 for the lymphatic markers (VEGFR3:YFP, LYVE1, PROX1, and PDPN in green, cyan, magenta, and white, respectively). The epididymis used for PLVAP detection is the contralateral organ of the one used for lymphatic markers. The white-squared regions in each photograph (left in caput and cauda) arc enlarged in the right photographs. PLVAP* micro-vascularization (red) is visible at the peritubular epithelium level of the initial segment (IS)/Sl and its expression markedly decreases in the other segments. At higher magnification, a network of blood capillaries surround each tubule mainly at the level of the IS/S1 while intcrtubular vessels can be observed in segment 3 (S3). In the cauda (S10), PLVAPpos vasculature is less dense and is mostly located in the intertubular space. The VEGFR3:YFP transgene (green), revealed by an anti-GFP antibody, is abundant in the caput and in particular in the 1S/S1. Enlargements show punctiform labeling at the peritubular level and in the intertubular zone. On can note the presence of very large lymphatics at the intertubular level (white star). In the cauda (S10), reactivity is mainly interstitial, stringy, and compatible with flattened lymphatic vessels trapped in the extracellular matrix. We also noted at the S9/S10 boundaries (delimited by the dotted line) the presence of large lymphatic vessels (white asterisk), suggesting that important drainage takes place at this location. The lymphatic marker LYVE1 in the caput is mainly interstitial and clearly stronger in the 1S/S1. We also noticed that the lymphatics detected with the transgcnc (VEGFR3:YFP) and with LYVE1 present differences (white arrows), in agreement with previous studies reporting heterogeneity for these markers at the level of the lymphatics (Pawlak and Caron, 2020; Ulvmar and Mäkinen, 2016). In the cauda, LYVE1 appears less present compared with VEGFR3:YFP. We also noticed differences in localization and intensity between these two lymphatic markers (white arrows). PROX1 (magenta) shows a profile comparable to those obtained for the transgcnc product and/or LYVE1 in both the caput and cauda. The labeling is punctiform (compatible with a nuclear localization) and filamentous at the level of the 1S/S1. The stringy appearance can be explained by the use of a PROX1-biotinylated antibody that generates a fuzzier/coarser signal. PDPN (white) appears abundant in the caput and cauda at the peritubular level of the epididymal epithelium and in the interstitial space. Stcrcocilia in the initial segment but not in other segments of the caput also show some reactivity with PDPN. Because of the fuzzy PDPN pattern, it is difficult at this resolution to assert that PDPN strictly colocalizes with VEGFR3:YFP, LYVE1 and PROX1. Panel B shows light-sheet confocal (12×) resolution of the caput/lS-S2 region of a VEGFR3 :YFP transgenic epididymis (green) or after PLVAP immunodetection (red). The superposition of the two channels (merge) suggests both co-localization but also specific territories. The far right picture shows the superposition of the surface rendering of the two markers, which allows distinguishing the lymphatic vasculature that expresses only the transgene (in dark green) from the PLVAP*/VEGFR3:YFP* vessels (in light green).

**Supplementary Figure 4:**
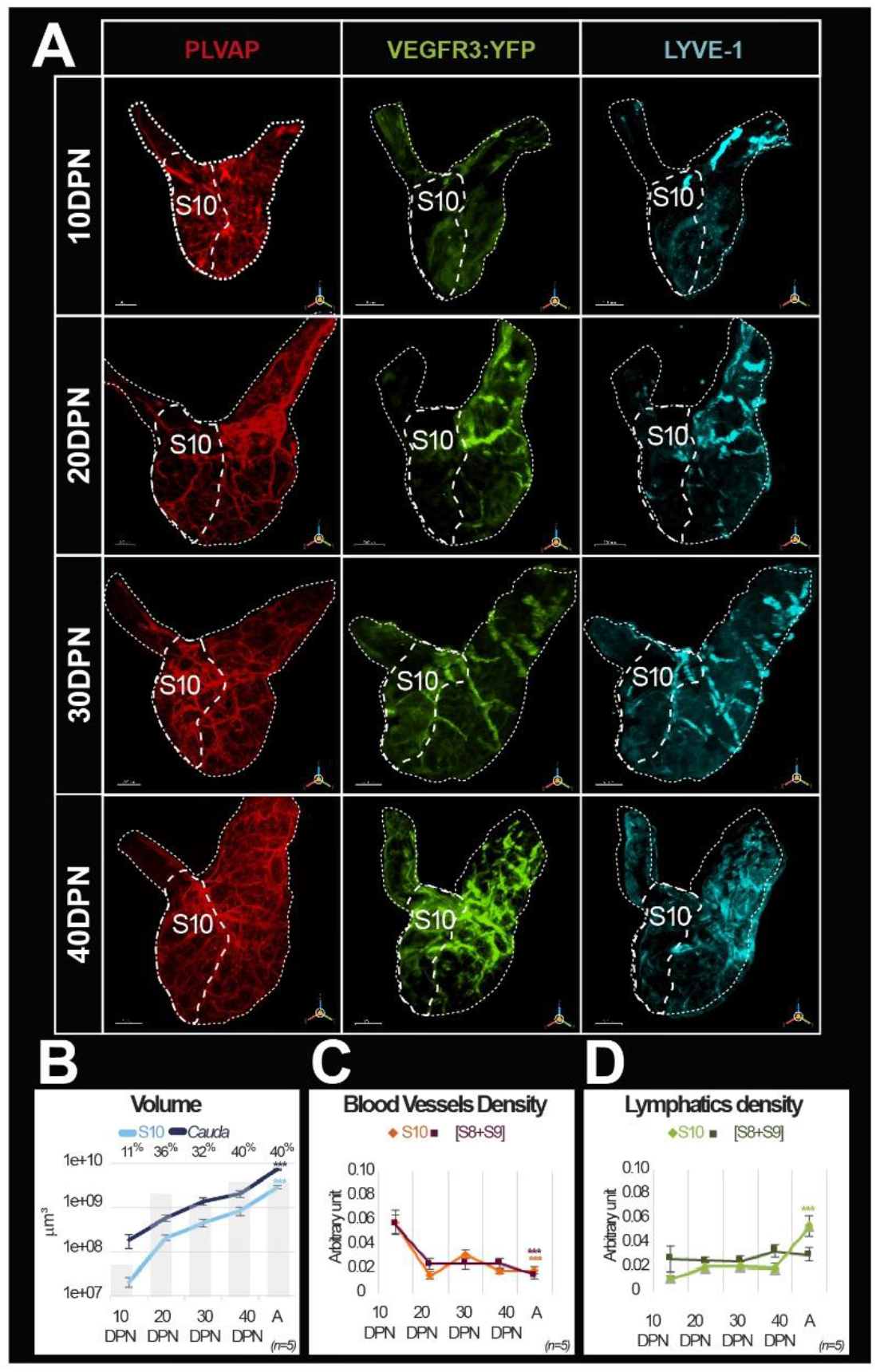
Evolution of the blood and lymphatic vascularization of the epididymal *cauda* during postnatal development. Panel A shows representative three-dimensional (3D) views of the blood and lymphatic networks at different postnatal stages of the *cauda* epididymis ontogenesis from 10 to 40 days postnatal (DPN). Whole-mount immunolabeling of PLVAP^1^ blood vessels (red) was conducted on the contralateral organ of the one used for the immunolabeling with the lymphatic marker LYVE1 (cyan) and detection of the YFP reporter gene (revealed using an anti-GFP antibody; green). The dotted line indicates the S8-S9/S10 border. The scale bars is 200 μm for 10 and 20 DPN, 300 μm for 30 DPN, and 400 μm for 40 DPN. Panel B shows evolution of the volume (in log 10) of the S10 segment (light blue line) and the *cauda* (S8 to S10; dark blue line) regions during postnatal development. The light grey histogram in the background gives the proportion of volume occupied by the S10 segment within the *cauda* at different postnatal developmental stages. Panels C and D present surface rendering of blood vessels (C) and lymphatics (D) evaluated using the IMARIS software as described above (see Figures 2 and 3). The curves represent the mean and standard error of the mean of the densities obtained for at least five individuals (for the postnatal development stages) and up to 13 individuals (for the adult stage). The color code in Panel C is orange =S10 and dark red =[S8+S9], In Panel D, the color code is light green =S10 and dark green = [S8+S9ļ. The Kruskal-Wallis test with Dunn’s post-test correction was used to determine statistical significance (*** p < 0.0001).

**Supplementary Figure 5:**
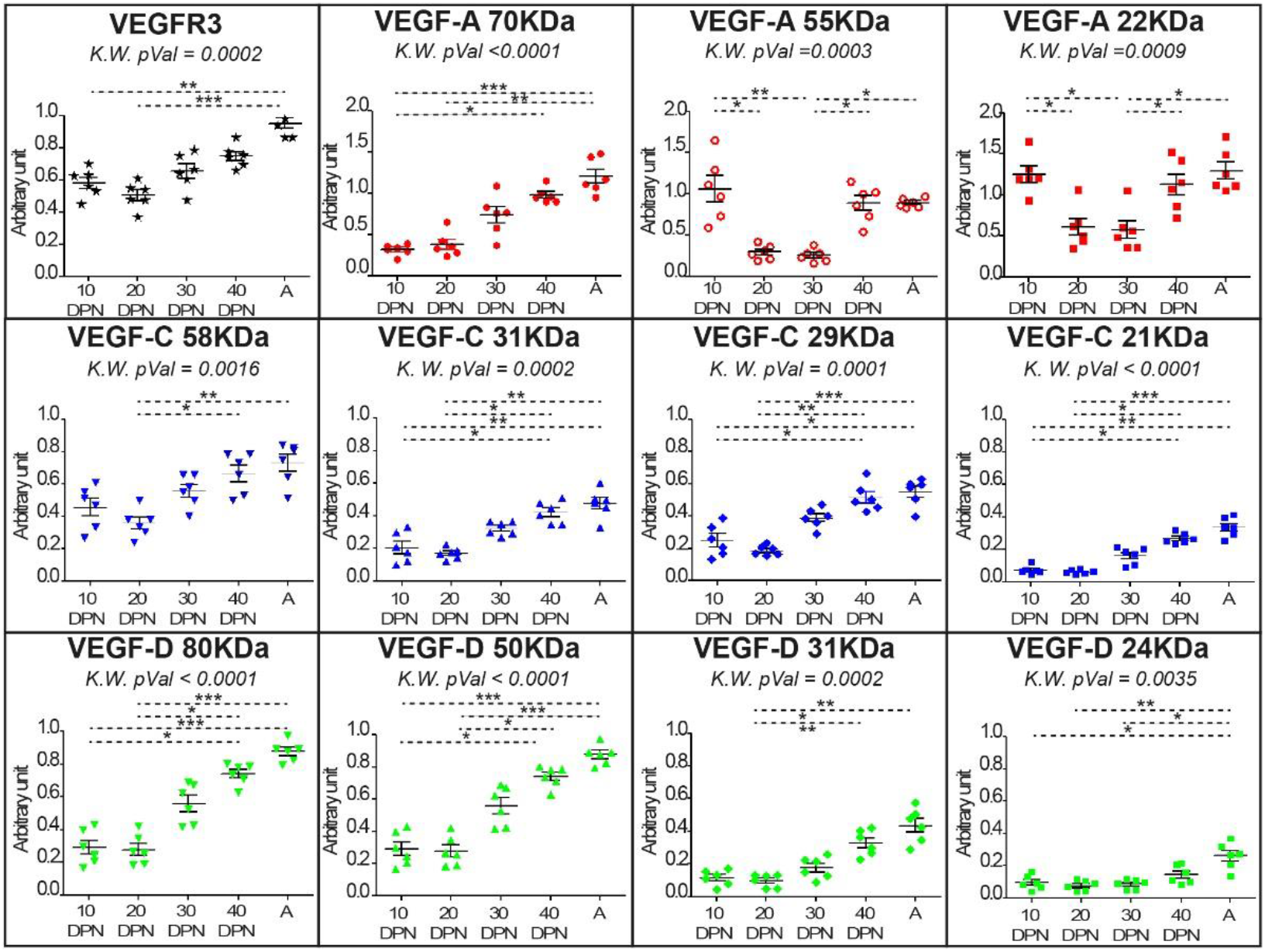
Quantification of all forms of the hemangiogenic and lymphangiogenic ligands VEGF-A, VEGF-C, and VEGF-D. Different forms of ligands during postnatal development of the epididymis were quantified by optical density measurement with ImageJ software. It was normalized by the level of GAPDH measured in the same organ. The profiles of VEGFR3 (black star) and the active isoforms of VEGF-A (22 kDa, red square), VEGF-C (21 kDa blue square), and VEGF-D (24 kDa, green square) arc those presented in Figure 5 and arc presented here for comparison with the other isoforms. We noted that both the 55 kDa and 22 kDa VEGF-A profiles behave similarly during postnatal development of the epididymis. There is a drop in these two forms at 20 days postnatal (DPN) followed by a plateau at 30 DPN and then a marked increase from 40 DPN onwards. VEGF-A could play a different role in the epididymis because the number of PLVAP^+^ blood vessels do not increase at these stages (sec Figure 4 and Supplementary Figure 4). Quantification of the different VEGF-C (blue) and VEGF-D (green) isoforms shows a pattern comparable to that observed for lymphatic density during postnatal development (see Figure 4 and Supplementary Figure 4). However, the present forms are mainly precursors, suggesting progressive lymphangiogenic activity.

**Supplementary Figure 6:**
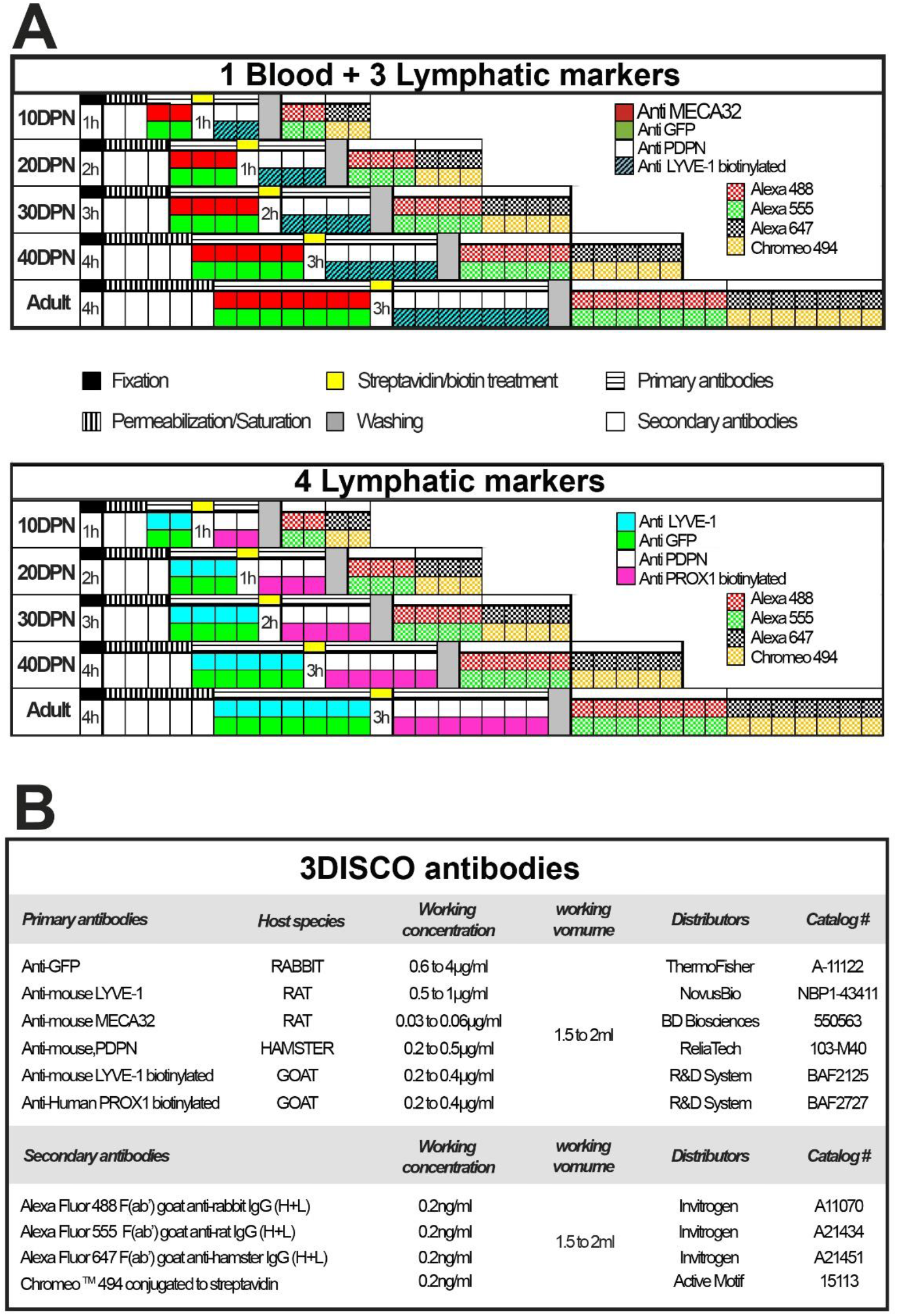
Multiplex labeling workflow used to visualize the blood and lymphatic vasculature of the mouse epididymis clarified by the 3DISCO method. Panel A presents the multiplex labeling schedule for blood and lymphatic immunodetection. Each square corresponds to 24 h except for the fixation and streptavidin/biotin treatment, where the time is indicated. Panel B provides detailed information regarding the various antibodies used in the course of the study.

**Supplementary Figure 7:**
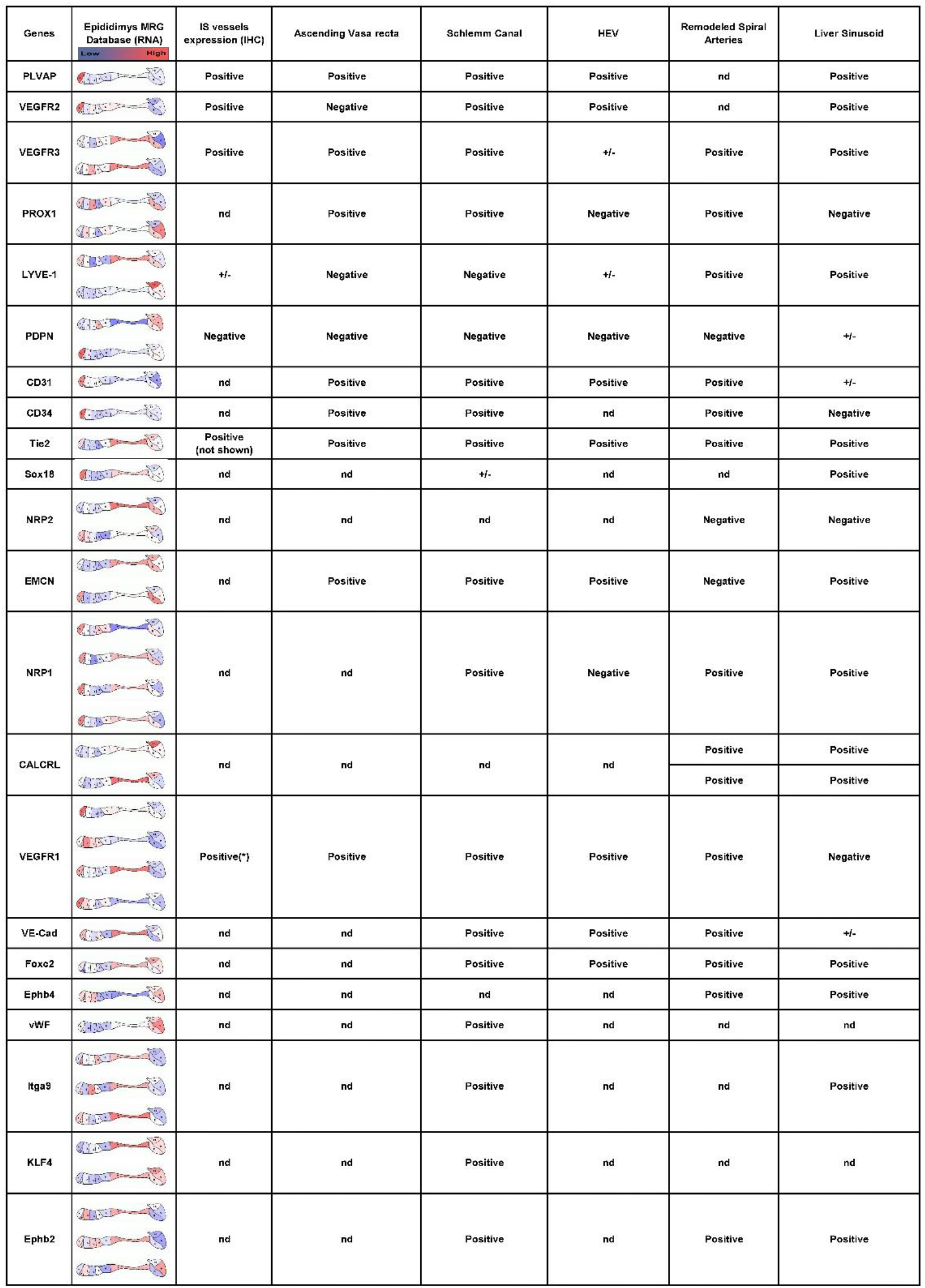
Expression of hybrid lymphatic markers in the epididymis. This lable compares markers associated with hybrid lymphaic vessels deseried in the literature (for a review, see pawlak et al., 2020) with (column 2) their expression a different segments of the epididymis (source Mammalian Repreduelive Genetic. www.mrgd.org). Multiple results for the same gene correspond to the use of different Oliyosels in the microurruys used by the MRG. Column 3 summurizes our present results at the initial segment level. The asterisk refers to the publication by Korpelainen et al. (1995).

Video1

https://ent.uca.fr/filez/x0ujshpf

Video2

https://ent.uca.fr/filez/3sjybraj5

Video3

https://ent.uca.fr/filez/o5ufb1

## Materials and Methods

### Mice

In this study, we have used 3–6-month-old male C57/B6 mice (Janvier Labs, France); the male VEGFR3-YFP mice came from the J. L. Thomas laboratory (UPMC-Inserm, Paris, France). All animals were housed according to institutional guideline with a 12-h photoperiod and food and water available *ad libitum*. Measures were taken to keep animal suffering to a minimum. The Auvergne Animal Experiment Ethics Committee (C2E2A) and the French Ministry approved all the following procedures for research (APAFIS authorization # 12376-2017112913113962 V3).

### Antibodies

The primary and secondary antibodies used in clearing methods are indicated in Supplementary Figure 6B. Additional antibodies used are: rat anti-mouse VEGFR3 (# NB110-61018, Novus Biologicals, LLC, Bio-Techne SAS, France); goat anti-mouse VEGF-A (# AF-493-SP, Novus Biologicals, LLC, Bio-Techne SAS, France), rabbit anti-mouse VEGF-C (# NB110-61022, Novus Biologicals, LLC, Bio-Techne SAS, France), goat anti-VEGF-D (# AF469, R&D Systems, Bio-Techne SAS, France), and rabbit anti-GAPDH (SAB2108668, Sigma Aldrich Chimie Sarl, France).

### Western blot analysis

Soluble epididymal proteins were prepared, as described previously (Chorfa et al., 2021), separated with sodium dodecyl sulfate–polyacrylamide gel electrophoresis (SDS-PAGE; 12% gel), and transferred to polyvinylidene fluoride (PVDF) membranes (Hybond ECL, Amersham Biosciences, Germany). Primary antibodies against the following protein were used: VEGFR3, VEGF-A, VEGF-C, VEGF-D, and GAPDH (served as a loading control). The appropriate HRP-conjugated secondary antibodies, goat anti-rabbit IgG or goat anti-mouse IgG (Abliance, France), were used to visualize the protein bands. Immunoreactive bands were detected by chemiluminescence (Clarity Western ECL Substrate Bio-Rad, France) using the ChemiDoc MP imaging system (Bio-Rad). Protein quantification was performed with ImageJ software. Protein amounts are expressed as relative values to the GAPDH.

### Whole-mount multiplex immunolabeling and clearing procedure

Epididymides were collected from mice after cervical dislocation and fixed in 4% paraformaldehyde (PAF) for a period of time determined by their size (see Supplementary Figure 6). Then, they were subjected to saturation/permeabilization for 1–5 days depending on their size in PBSGTS solution (1X saline buffer phosphate containing 2% gelatin, 0.5% triton X100, and saponin 1 μg/ml) on a rotary shaker (100 rpm) at room temperature. Incubation with primary and secondary antibodies was performed under rotation at 37°C in the same buffer according to the schedule described in Supplementary Figure 6A. Concentrations of primary and secondary antibodies used are reported in Supplementary Figure 6B. The immunolabeled epididymides were then embedded in a 1.5% agarose cube in 0.5x Tris–Acetate– EDTA buffer and cleared according the 3DISCO method (Belle et al., 2014). The sample were stored in a dark place at room temperature until observation.

### Immunofluorescence

Paraffin-embedded 5 μm sections of epididymides were subjected to heat-induced antigen recovery and then permeabilized for 30 min at room temperature with PBS supplemented with 0.3% Tween 20 and saponin (1 mg/mL). The sections were incubated for 1 h at room temperature in the same solution with 10% serum before overnight incubation with the primary antibody at 4°C. Their detection was then carried out with appropriate secondary antibodies conjugated to Alexa Fluor 488, Alexa Fluor 555, or Alexa Fluor 647 (Invitrogen, Thermofisher Scientific, France). Nuclei were stained with Hoechst solution (1 μg/μL). The sections were embedded in Mowiol 4-88 (Sigma Aldrich Chimie Sarl, France) and stored at 4°C until observation.

### Image acquisition

#### Macroscopic microscopy

Macroscopic observations of endogenous VEGFR3:YFP transgene expression of the epididymis and testis were performed using with a Leica binocular magnifier (with a 1× objective).

#### Confocal microscopy

Multiplex immunofluorescence acquisition was performed using a SP8 confocal laser scanning microscope equipped with a Plan Apo λ 40X Oil objective (Leica, Germany). The following parameters were used: pinhole size of 1 airy unit (AU); Z-step: 0.5 μm. Four acquisition sequences were executed: (1) for Alexa 647 detection (638 nm laser 659–698 nm window, using a photomultiplier tubes (PMT) detector at 700 V gain); (2) for Alexa 488 detection (488 nm laser; 493–541 nm window, using a PMT detector at 565 V gain) and for Chromeo 494 detection (488 nm laser; 660–700 nm window, using a PMT detector with a gain of 800 V); (3) for Alexa 555 detection (laser 552 nm; 560–570 nm window, using a PMT detector with a gain of 700 V); and (4) for Hoechst detection (laser 405 nm; 423–518 nm window, using a Hybrid detector with 10% gain).

#### Light-sheet ultramicroscopy

3D imaging was achieved with an ultramicroscope (LaVision BioTec Miltenyi, Germany) using ImspectorPro software. The images were obtained with either a MI PLAN *2x*/NA 0.5, a MI PLAN 4X/0.35 objectives (MVPLAPO, Olympus), or a 20x/0.95 objective (Leica). Each sample was placed in a 100% quartz imaging reservoir filled with dibenzyl ether (DBE) and illuminated from the side by the laser sheet. Images were acquired with a Andor Neo SCMOS CCD camera (2160 × 2160 pixels, LaVision BioTec). All 3D acquisitions were performed with a step size between each image was fixed at 1 μm (1.6 μm/pixel). Lasers at 488, 561, and 635 nm were used to obtained images using two light sources for the *caput, corpus*, or *cauda* epididymis.

### Image processing

Ultramicroscopy data sets were uploaded to IMARIS version 9.7 (Bitplane, Oxford Instruments England). The stacks were converted to IMARIS files (.ims) and 3D visualizations of z-stack images were generated using the volume rendering function. The vessels were transformed into surface rendering and the volume calculated automatically by IMARIS. Vessel densities were estimated as the ratio of the vessel volume to the total volume of the region. The videos were made IMARIS without deconvolution of the image.

### Statistical analysis

All experiments were repeated at least three times and representative images are shown. All statistical analyses were performed with Prism software (GraphPad, USA). The Mann–Whitney test was used to compare two groups. The Kruskal–Wallis test followed by Dunn’ s post-test was used to compare more than two groups. A two-way analysis of variance (ANOVA) test with a Bonferroni post-test was used to analyze the covariance of ligands (VEGF-A, VEGF-C, and VEGF-D) with the VEGFR3 receptor. The co-localization of the ligands VEGF-C and VEGF-D was evaluated by Pearson correlation and Mander’s overlap coefficient with IMARIS co-localization modules (Bitplane). The values for colocalization analysis represent the mean ± standard error of the mean (SEM).

## Acknowledgements

The authors would like to thank the CLIC confocal microscopy platform (GReD Institute, Clermont-Ferrand, France) for their expert assistance during this study.

## Authors contribution

JHB initiated the project, designed and carried out all experiments with the help of CG, SB, NP, and CV. JHB, CD, and AC tested various organ clearing protocols with contributions from CG and SB. JHB and AB performed the image analysis and recorded the videos. JLT provided the transgenic mice used in this study. AB and MT performed the development of the light-sheet microscopy technology after organ clearing. LP, LPF, RG, FS, AK, AB, and MT provided valuable comments for data analysis and interpretation. JHB and JRD wrote the paper.

